# β-RA targets mitochondrial metabolism and adipogenesis, leading to therapeutic benefits against CoQ deficiency and age-related overweight

**DOI:** 10.1101/2021.04.13.438670

**Authors:** Agustín Hidalgo-Gutiérrez, Eliana Barriocanal-Casado, María Elena Díaz-Casado, Pilar González-García, Riccardo Zenezini Chiozzi, Darío Acuña-Castroviejo, Luis Carlos López

## Abstract

Primary mitochondrial diseases are caused by mutations in mitochondrial or nuclear genes, leading to abnormal function of specific mitochondrial pathways. Mitochondrial dysfunction is also a secondary event in more common pathophysiological conditions, such as obesity and metabolic syndrome. In both cases, the improvement and management of mitochondrial homeostasis remains challenging. Here, we show that beta-resorcylic acid (β-RA), a natural phenolic compound, competes *in vivo* with 4-hydroxybenzoic acid, the natural precursor of Coenzyme Q biosynthesis. This leads to a decrease of demethoxyubiquinone, an intermediate metabolite of CoQ biosynthesis that is abnormally accumulated in *Coq9*^*R239X*^ mice. As a consequence, β-RA rescues the phenotype of *Coq9*^*R239X*^ mice, a model of primary mitochondrial encephalopathy. Moreover, we observe that long-term treatment with β-RA also reduces the size and content of the white adipose tissue (WAT) that is normally accumulated during aging in wild-type mice, leading to a prevention of hepatic steatosis and an increase in survival at the old stage of life. The reduction in WAT content is due to a decrease in adipogenesis, an adaptation of the mitochondrial proteome in the kidneys, and a stimulation of glycolysis and acetyl-CoA metabolism. Therefore, our results demonstrate that β-RA acts through different cellular mechanisms, with effects on mitochondrial metabolism, and it may be used for the treatment of primary Coenzyme Q deficiency, overweight, and hepatic steatosis.

## Introduction

Mitochondria are the primary sites of cellular energy production and have also a broad range of metabolic functions. Thus, mitochondrial dysfunction can produce far-ranging, varied, and severe consequences. Mitochondrial dysfunction can be directly caused by mutations in mitochondrial DNA or mutations in nuclear genes that encode mitochondrial proteins, leading to primary mitochondrial diseases. Aside from direct causes, mitochondrial dysfunction can also occur as a secondary event in more common diseases, such as neurodegenerative diseases, obesity, or metabolic syndrome.

One particular case of mitochondrial disease is Coenzyme Q10 (CoQ_10_) deficiency syndrome, which can be primarily caused by mutations in genes that encode proteins involved in the CoQ_10_ biosynthetic pathway (primary CoQ_10_ deficiency). Primary CoQ_10_ deficiency presents heterogeneous clinical phenotypes depending on the specific mutation in the CoQ biosynthesis pathway (Desbats, Morbidoni et al., 2016, Luna-Sanchez, Diaz-Casado et al., 2015). But also, especially given the variety of functions of CoQ, multiple pathomechanisms are induced by low levels of CoQ, including declined bioenergetics (Garcia-Corzo, Luna-Sanchez et al., 2013, Lopez, Quinzii et al., 2010, Luna-Sanchez et al., 2015, Quinzii, Lopez et al., 2010, Quinzii, Lopez et al., 2008b), increased oxidative stress (Duberley, Abramov et al., 2013, Quinzii, Garone et al., 2013, Quinzii et al., 2010, Quinzii et al., 2008b), disrupted sulfide metabolism (Kleiner, Barca et al., 2018, Luna-Sanchez, Hidalgo-Gutierrez et al., 2017), and defective *de novo* pyrimidine biosynthesis (Lopez-Martin, Salviati et al., 2007).

CoQ_10_ deficiency can also be induced as a secondary effect of certain drugs (Mancuso, Orsucci et al., 2010) and triggered indirectly by other diseases, including multifactorial diseases and disorders caused by mutations in genes not related to the CoQ_10_ biosynthesis pathways (Emmanuele, Lopez et al., 2012, Quinzii, Lopez et al., 2008a, Turunen, Olsson et al., 2004, Yubero, Montero et al., 2016). Metabolic syndrome is a multifactorial disease with secondary mitochondrial dysfunction. The white adipose tissue (WAT) and skeletal muscle from patients and mice with insulin resistance, a characteristic usually associated to metabolic syndrome, show decreased levels of the CoQ biosynthetic proteins COQ7 and COQ9, leading to reduced CoQ levels in the mitochondria (Fazakerley, Chaudhuri et al., 2018).

In experimental cases of CoQ_10_ deficiency, the levels of CoQ_10_ in blood, cells and tissues could be increased by exogenous CoQ_10_ supplementation. However, CoQ_10_ has very low absorption and bioavailability when it is orally administrated, and a very low proportion of this exogenous CoQ_10_ can reach the mitochondria of the cells in most tissues (Bentinger, Dallner et al., 2003, Garcia-Corzo, Luna-Sanchez et al., 2014). Thus, Hydroxybenzoic acid derivatives (HBAs) have been proposed as an alternative strategy to attenuate CoQ_10_ deficiency since they have been shown to modulate the endogenous CoQ biosynthetic pathway (Herebian, Lopez et al., 2018). HBAs constitute a group of natural phenolic compounds present in plants with a general structure of the C6–C1 type derived from benzoic acid. Variable positioning of hydroxyl and methoxy groups on the aromatic ring produce several different compounds, like 2-hydroxybenzoic acid (or salicylic acid), 4-hydroxybenzoic acid (4-HB), 2,4-dihydroxybenzoic acid (2,4-diHB; or β-resorcylic acid; β-RA), and 4-hydroxy-3-methoxybenzoic acid (or vanillic acid; VA). Interestingly, β-RA has the hydroxyl group that is incorporated to the benzoic ring during CoQ biosynthesis. This hydroxylation step is catalyzed by COQ7, which uses demethoxyubiquinone (DMQ) as a substrate and requires the COQ9 protein for its normal function and stability (Garcia-Corzo et al., 2013). As a consequence, the administration of high doses of β-RA bypasses the defects in the COQ7 reaction, leading to a dramatic increase in the survival of *Coq7* conditional knockout mice and the *Coq9*^*R239X*^ mice due to increased levels of CoQ and/or to decreased levels of DMQ in the kidneys, heart, skeletal muscle and intestine (Hidalgo-Gutierrez, Barriocanal-Casado et al., 2019, Wang, Oxer et al., 2015, Wang, Smith et al., 2017). In *Coq9*^*R239X*^ mice, those biochemical changes resulted in significant improvements of encephalopathic features like astrogliosis and spongiosis (Hidalgo-Gutierrez et al., 2019). Similarly, the supplementation with high doses of β-RA to podocyte-specific *Coq6* or *Adck4* (*Coq8b*) knockout mice prevented renal dysfunction and increased survival, although the effect of β-RA on CoQ metabolism in these mouse models was not reported and, therefore, the therapeutic mechanisms of those cases are unknown (Widmeier, Airik et al., 2019a, Widmeier, Yu et al., 2019b). Additionally, Wang and colleagues reported that β-RA decreased the body weight of wild-type mice and increased survival in animals at the middle-age and old stages of life, but the mechanisms behind these observations remain to be elucidated. Consequently, these results in the *Coq6* and *Adck4* mouse models and in wild-type mice suggest that β-RA may work through additional unidentified mechanisms.

Here, we have evaluated whether a lower dose of β-RA, which may increase its translational potentiality, leads to therapeutic outcomes in the encephalopathic *Coq9*^*R239X*^ mice and if that effect is mainly due to β-RA interference in CoQ metabolism. Additionally, we have tested whether β-RA could be a useful agent to treat the fat accumulation linked to aging.

## Results

### β-RA induces phenotypic and morphological benefits against both age-related obesity and mitochondrial encephalopathy due to CoQ deficiency

β-RA was incorporated into the chow of both wild-type and *Coq9*^*R239X*^ mice at a concentration of 0.33 % (w/w), which gives a dose of 0.4-0.7 g/kg b.w./day, considering the animal food intake, which was similar in all groups (Fig. 1A-C). This low dose of β-RA improved the survival of *Coq9*^*+/+*^ mice at the old stage of life (Fig. 1D-E), being alive the 87% of the treated *Coq9*^*+/+*^ mice against a 62% of untreated mice. However, the survival curve becomes similar to the survival curve of untreated animals after 28 months of age. Similarly, the low dose treatment of β-RA also improved the survival of *Coq9*^*R239X*^ mice (Fig. 1D), and we even observed a maximal lifespan higher than the maximal lifespan reported when *Coq9*^*R239X*^ mice were treated with a high dose of β-RA (Hidalgo-Gutierrez et al., 2019).

**Figure 1.**
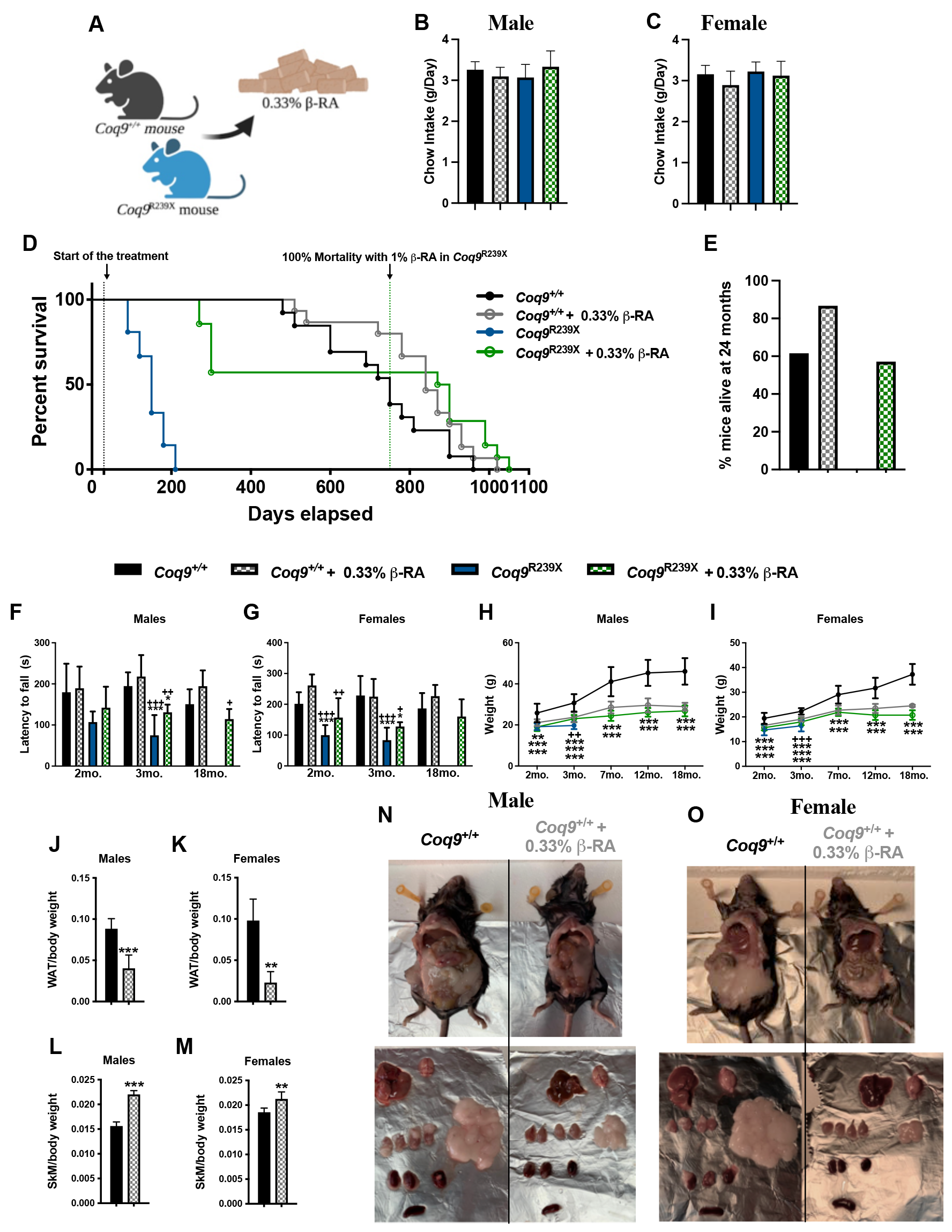
Survival and phenotypic characterization of *Coq9*^*+/+*^ and *Coq9*^*R239X*^ mice during the supplementation with 0.33% β-RA. (**A**)Schematic figure of the β β-RA treatment in *Coq9*^+/+^ and *Coq9*^R239X^ mice. (**B** and **C**) Daily food intake in male and female *Coq9*^+/+^ and *Coq9*^R239X^ mice. (**D**) Survival curve of *Coq9*^*+/+*^ mice, *Coq9*^*+/+*^ mice under 0.33% β -RA supplementation, *Coq9*^*R239X*^ mice, and *Coq9*^*R239X*^ mice under 0.33% β -RA supplementation. The treatments started at 1 month of age [Log-rank (Mantel-Cox) test or Gehan–Breslow–Wilcoxon test; *Coq9*^+/+^ mice, n = 13; *Coq9*^*+/+*^ mice under 0.33% β-RA supplementation, n = 15; *Coq9*^*R239X*^ mice, n = 21; *Coq9*^*R239X*^ mice under 0.33% β-RA supplementation, n = 14. (**E**) Percentage of mice alive at 24 months of age. (**F** and **G**) Rotarod test of male and female *Coq9*^*+/+*^ mice, *Coq9*^*+/+*^ mice under 0.33% β-RA supplementation, *Coq9*^*R239X*^ mice, and *Coq9*^*R239X*^ mice under 0.33% β-RA supplementation. (**H** and **I**) Body weight of male and female *Coq9*^*+/+*^ mice, *Coq9*^*+/+*^ mice under 0.33% β-RA supplementation, *Coq9*^*R239X*^ mice, and *Coq9*^*R239X*^ mice under 0.33% β-RA supplementation. (**J** to **M**) Weight of the epididymal, mesenteric and inguinal white adipose tissue (WAT) (J and K) and hind legs skeletal muscle (SKM) (L and M) relative to the total body weight in male and female *Coq9*^*+/+*^ mice, and *Coq9*^*+/+*^ mice under 0.33% β-RA supplementation at 18 months of age. (**N** and **O**) Representative images of male (N) and female O) mice, and their tissues, at 18 months of age, both untreated and treated. Data are expressed as mean ± SD. *P < 0.05, **P < 0.01, ***P < 0.001, differences *versus Coq9*^*+/+*^; +P < 0.05, ++P < 0.01, +++P < 0.001, *Coq9*^*+/+*^ mice under 0.33% β-RA supplementation; (one-way ANOVA with a Tukey’s post hoc test or t-test; n = 5–34 for each group).

The encephalopathic features of *Coq9*^*R239X*^ mice result in characteristics of lower locomotor activity and uncoordination. However, *Coq9*^*R239X*^ mice improved after β-RA administration compared to untreated *Coq9*^*R239X*^ mice. The treatment did not significantly affect the results of the rotarod test in wild-type animals (Fig. 1F-G). Both *Coq9*^*+/+*^ and *Coq9*^*R239X*^ mice treated with β-RA had a healthy appearance (Movies S1 and S2).

The body weights were significantly reduced in both male and female *Coq9*^*+/+*^ mice after one month of treatment, reaching a maximal weight of about 28 grams in males and 23 grams in females at seven months of age. These weights were then maintained throughout the remaining life of the animals (Fig. 1H-I) (Movie S3). Curiously, the treatment with β-RA slightly increased the body weights of *Coq9*^*R239X*^ mice, which usually weigh less than their untreated *Coq9*^*+/+*^ littermates (Fig. 1H-I). Consequently, both treated *Coq9*^*+/+*^ and treated *Coq9*^*R239X*^ mice had similar body weights. The reduced body weight in *Coq9*^*+/+*^ mice after β-RA treatment was mainly caused by the prevention of accumulation of WAT (Fig. 1J,K,N,O) while still preserving the content, weight, and strength of the skeletal muscle (Fig. 1L-O; Fig. S1).

The most notable histopathological features of CoQ_10_ deficiency in *Coq9*^*R239X*^ mice is cerebral spongiosis and astrogliosis (Fig. 2A1-D1). Low-dose β-RA supplementation in *Coq9*^*R239X*^ mice for two months decreased the characteristic spongiosis (Fig. 2E1-F1) and astrogliosis (Fig. 2G1-H1), similar to the therapeutic effect previously reported with a higher dose (Hidalgo-Gutierrez et al., 2019). In *Coq9*^*+/+*^ mice, β-RA supplementation for two months did not produce significant morphological alterations in the brain (Fig. 2I1-P1), liver (Fig. S2A-F), kidneys (Fig. S2G-L), spleen (Fig. S2M-P), heart (Fig. S2Q-T), or small intestine (Fig. S2U-X), and the blood and urine markers of renal and hepatic functions did not reveal any abnormality (Table S1).

**Figure 2.**
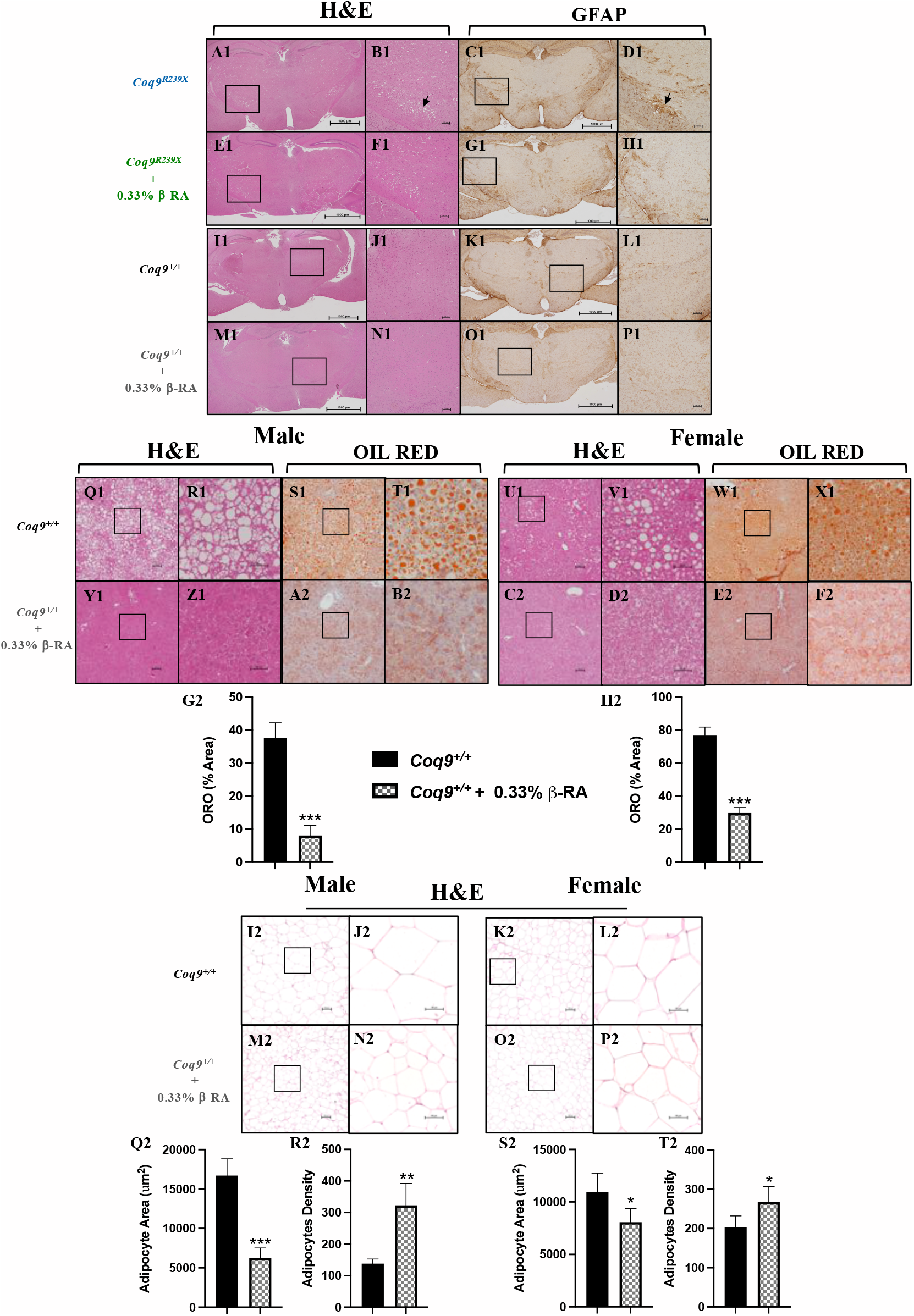
Morphological evaluation of symptomatic tissues from *Coq9*^*R239X*^ and *Coq9*^*+/+*^ mice under the supplementation with 0.33% β-RA. (**A1 to P1)** H&E stain and anti-GFAP immunohistochemistry in sections of the diencephalon from *Coq9*^*R239X*^ mice (A1 to D1), *Coq9*^*R239X*^ mice under 0.33% of β-RA supplementation (E1 to H1), *Coq9*^*+/+*^ mice (I1 to L1), *Coq9*^*+/+*^ mice under 0.33% β-RA supplementation (M1 to P1) at 3 months of age. (**Q1 to F2**) H&E and oil-red stains in sections of the liver from male (Q1 to T1) and female (U1 to X1) Coq9^+/+^ mice and male (Y1 to B2) and female (C2 to F2) *Coq9*^*+/+*^ mice under 0.33% β-RA supplementation at 18 months of age. (**G2 and H2**) Percentage of the area corresponding to the Oil Red O stains in sections of the liver from *Coq9*^*+/+*^ mice and *Coq9*^*+/+*^ mice under 0.33% β-RA supplementation at 18 months of age. (**I2 to P2**) H&E stain in sections of the epididymal WAT from male (G2-H2) and female (I2-J2) *Coq9*^*+/+*^ mice and male (K2-L2) and female (M2-N2) *Coq9*^*+/+*^ mice under 0.33% β-RA supplementation at 18 months of age. (**Q2 to T2**) Average of the area of each adipocyte and the adipocytes density in sections of the epididymal WAT from *Coq9*^*+/+*^ mice and *Coq9*^*+/+*^ mice under 0.33% β-RA supplementation at 18 months of age. Data are expressed as mean ± SD. *P < 0.05, **P < 0.01, ***P < 0.001, differences *versus Coq9*^*+/+*^(t-test; n = 4–6 for each group).

At 18 months of age, the liver of both male and female wild-type mice showed features of steatosis (Fig. 2Q1-X1 and G2-H2). Chronic supplementation with β-RA dramatically reduced the signs of hepatic steatosis (Fig. 2Y1-F2 and G2-H2). Non-alcoholic hepatic steatosis is frequently associated with fat accumulation. Consequently, epididymal WAT showed characteristics of hypertrophy in both male and female *Coq9*^*+/+*^ mice at 18 months of age (Fig. 2I2-L2 and Q2-T2), with adipocytes that were bigger in size and lower in number per area. β-RA supplementation suppressed the epididymal WAT hypertrophy in both male and female *Coq9*^*+/+*^ mice at 18 months of age (Fig. 2M2-P2 and Q2-T2).

### β-RA leads to bioenergetics improvement in *Coq9*^*R239X*^ mice through its direct participation in the CoQ biosynthetic pathway

The decrease of DMQ_9_ was previously reported as the main therapeutic mechanism of a high-dose of β-RA in the treatment in *Coq9*^*R239X*^ mice, although the effects in the CoQ biosynthetic pathway in wild-type animals were not evaluated (Hidalgo-Gutierrez et al., 2019). Thus, we have evaluated whether a lower dose of β-RA interferes with CoQ biosynthesis in both *Coq9*^*+/+*^ and *Coq9*^*R239X*^ mice. In *Coq9*^*+/+*^ mice, β-RA induced very mild changes in the tissue levels of CoQ_9_, CoQ_10_, and DMQ_9_ (Fig. 3A1-L1, Fig. S4A-D and Fig. S5A-B). The levels of CoQ_9_ were similar in the brain, kidneys, liver heart and WAT of untreated and treated wild-type mice, whilst in skeletal muscle β-RA induced a mild reduction in the levels of CoQ_9_ (Fig. 3A1-D1, Fig. S4A and Fig. S5A). DMQ_9_ was undetectable in the tissues of untreated wild-type mice, and β-RA supplementation induced the accumulation of very low levels of DMQ_9_ in the kidneys, liver, skeletal muscle and WAT but not in the brain or heart (Fig. 3I1-L1, Fig. S4C and Fig. S5B). Consequently, the ratio DMQ_9_/CoQ_9_ was not significantly altered in *Coq9*^*+/+*^ mice treated with β-RA, as it is observed in the untreated *Coq9*^*R239X*^ mice (Fig. 3M1-P1). In *Coq9*^*R239X*^ mice, β-RA administration induced a mild increase of CoQ_9_ in the kidneys (Fig. 3B1 and Fig. S3B) compared to untreated *Coq9*^*R239X*^ mice. However, the levels of CoQ_9_ did not change in the brain, liver, skeletal muscle, or heart of *Coq9*^*R239X*^ mice after the β-RA treatment (Fig. 3A1,C1,D1 and Fig. S4A). Remarkably, the levels of DMQ_9_ and, consequently, the DMQ_9_/CoQ_9_ ratio, were significantly decreased in the kidneys (Fig. 3J1,N1 and Fig. S3B), liver (Fig. 3K1,O1), skeletal muscle (Fig. 3L1,P1), and heart (Fig. S4C,D) of the *Coq9*^*R239X*^ mice treated with β-RA compared to untreated *Coq9*^*R239X*^ mice. However, β-RA did not reduce the levels of DMQ_9_ or the DMQ_9_/CoQ_9_ ratio in the brain of the *Coq9*^*R239X*^ mice (Fig. 3I1,M1), as it was also reported in the treatment with the higher dose of β-RA (Hidalgo-Gutierrez et al., 2019). Therefore, the effect of β-RA on CoQ metabolism in *Coq9*^*R239X*^ mice in this study is similar to the effect previously reported with a higher dose of β-RA (Hidalgo-Gutierrez et al., 2019).

**Figure 3.**
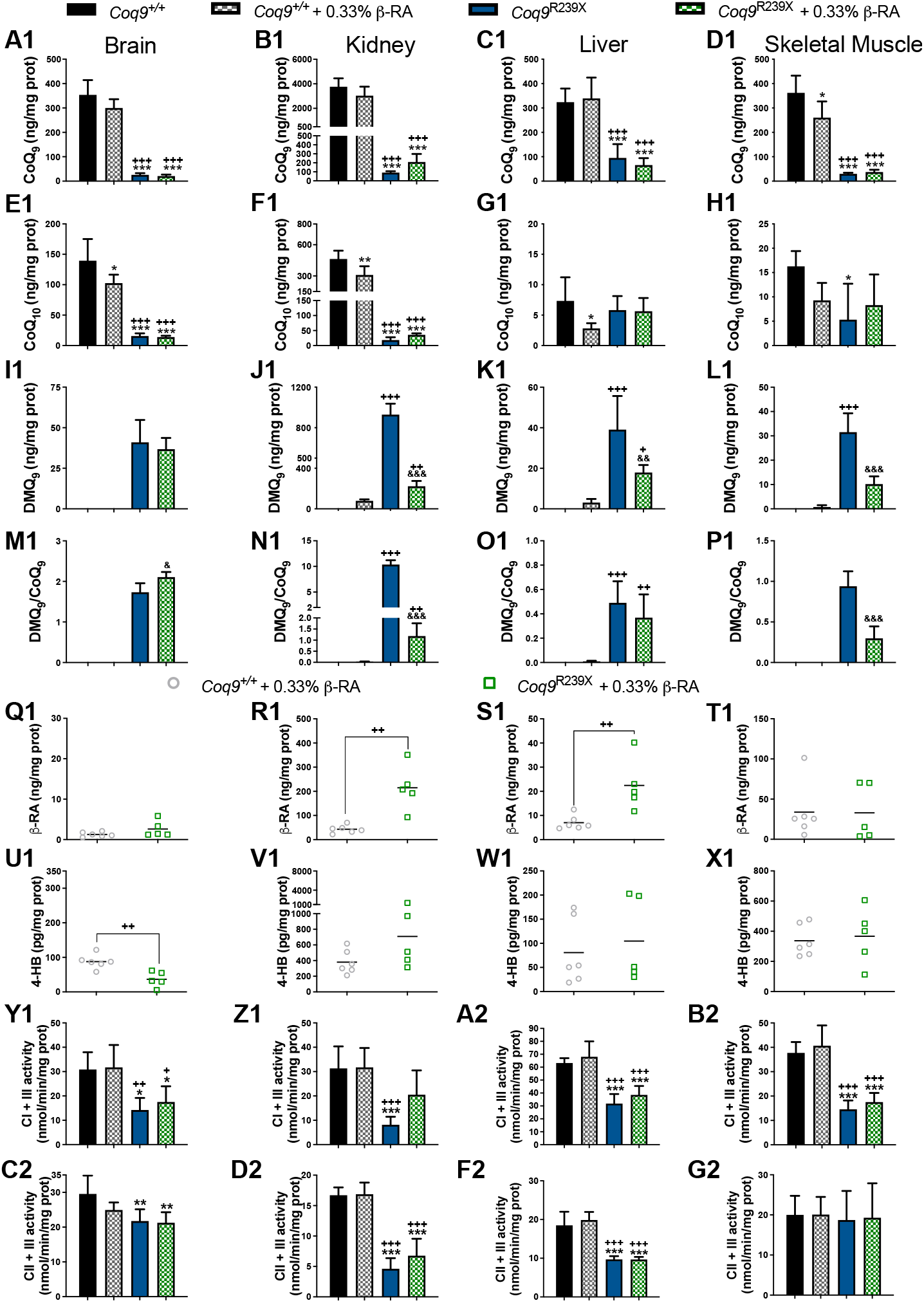
CoQ metabolism and mitochondrial function in the tissues from *Coq9*^*+/+*^ mice, *Coq9*^*+/+*^ mice under the supplementation with 0.33% β-RA, *Coq9*^*R239X*^ mice and *Coq9*^*R239X*^ mice under the supplementation with 0.33% β-RA. (**A1** to **D1**) Levels of CoQ9 in the brain (A1), kidneys (B1), liver (C1) and hind legs skeletal muscle (D1) from *Coq9*^*+/+*^ mice, *Coq9*^*+/+*^ mice under 0.33% β-RA treatment, *Coq9*^*R239X*^ mice, and *Coq9*^*R239X*^ mice under 0.33% β-RA treatment. (**E1** to **H1**) Levels of CoQ10 in the brain (E1), kidneys (F1), liver (G1) and hind legs skeletal muscle (H1) from *Coq9*^*+/+*^ mice, *Coq9*^*+/+*^ mice under 0.33% β-RA treatment, *Coq9*^*R239X*^ mice, and *Coq9*^*R239X*^ mice under 0.33% β-RA treatment. (**I1** to **L1**) Levels of DMQ_9_ in the brain (I1), kidneys (J1), liver (K1) and hind legs skeletal muscle (L1) from *Coq9*^*+/+*^ mice, *Coq9*^*+/+*^ mice under 0.33% β-RA treatment, *Coq9*^*R239X*^ mice, and *Coq9*^*R239X*^ mice under 0.33% β-RA treatment. Note that DMQ9 is not detected in samples from *Coq9*^*+/+*^ mice. (**M1** to **P1**) DMQ9/CoQ9 ratio in the brain (M1), kidneys (N1), liver (O1) and hind legs skeletal muscle (P1) from *Coq9*^*+/+*^ mice, *Coq9*^*+/+*^ mice under 0.33% β-RA treatment, *Coq9*^*R239X*^ mice, and *Coq9*^*R239X*^ mice under 0.33% β-RA treatment. (**Q1** to **X1**) Levels of β-RA in the brain (Q1), kidneys (R1), liver (S1) and hind legs skeletal muscle (T1) from *Coq9*^*+/+*^ mice under 0.33% β-RA treatment and *Coq9*^*R239X*^ mice under 0.33% β-RA treatment. β-RA were undetectable in *Coq9*^*+/+*^ mice and *Coq9*^*R239X*^ mice. (**U1** to **X1**) Levels of 4-HB in the brain (U1), kidneys (V1), liver (W1) and hind legs skeletal muscle (X1) from *Coq9*^*+/+*^ mice under 0.33% β-RA treatment and *Coq9*^*R239X*^ mice under 0.33% β-RA treatment. (**Y1** to **B2**) Complex I + III (CI + III) activities in the brain (Y1), kidneys (Z1), liver (A2) and hind legs skeletal muscle (B2) from *Coq9*^*+/+*^ mice, *Coq9*^*+/+*^ mice under 0.33% β-RA treatment, *Coq9*^*R239X*^ mice, and *Coq9*^*R239X*^ mice under 0.33% β-RA treatment. (**C2** to **G2)** Complex II+ III (CII + III) activities in the brain (C2), kidneys (D2), liver (F2) and hind legs skeletal muscle (G2) from *Coq9*^*+/+*^ mice, *Coq9*^*+/+*^ mice under 0.33% β-RA treatment, *Coq9*^*R239X*^ mice, and *Coq9*^*R239X*^ mice under 0.33% β-RA treatment. Tissues from mice at 3 months of age. Data are expressed as mean ± SD. *P < 0.05; **P < 0.01; ***P < 0.001; differences versus *Coq9*^*+/+*^. +P < 0.05; ++P < 0.01; +++P < 0.001, differences *versus Coq9*^+/+^ under 0.33% β-RA treatment. &P < 0.05; &&P < 0.01; &&&P < 0.001; differences *versus Coq9^R239X^*. (one-way ANOVA with a Tukey’s post hoc test or t-test; n = 5–8 for each group).

The tissue-specific reduction in the levels of DMQ_9_ in *Coq9*^*R239*X^ mice seemed to correlate with the increase of β-RA, since the levels of β-RA were higher in the kidneys (Fig. 3R1), liver (Fig. 3S1), skeletal muscle (Fig. 3T1), and heart (Fig. S4E) than in the brain (Fig. 3Q1) of *Coq9*^*R239*X^ mice. The levels of 4-HB, the natural precursor for CoQ biosynthesis, did not increase in response to the treatment with β-RA in any tissue of either *Coq9*^*+/+*^ or *Coq9*^*R239X*^ mice (Fig. 3U1-X1; Fig. S4F).

Bioenergetically, the treatment with β-RA did not produce any changes in the brain in either *Coq9*^*+/+*^ or *Coq9*^*R239X*^ mice (Fig. 3Y1, C2 and S6A), but it did increase the activities of complexes I + III and II + III (Fig. 3Z1,D2) and mitochondrial respiration (Fig. S6B) in the kidneys of the treated *Coq9*^*R239X*^ mice compared to the untreated *Coq9*^*R239X*^ mice. These data are comparable to those reported for the treatment with the high dose of β-RA (Hidalgo-Gutierrez et al., 2019). Other tissues did not experience major changes in mitochondrial bioenergetics in *Coq9*^*+/+*^ or *Coq9*^*R239X*^ mice (Fig. 3Y1-G2; Fig. S4G-H).

Because β-RA is an analog of 4-HB, its effects in reducing DMQ_9_ in *Coq9*^*R239X*^ mice is most likely due to its competition with 4-HB in entering the CoQ biosynthetic pathway through the activity of COQ2. To investigate this hypothesis, we supplemented *Coq9*^*+/+*^ and *Coq9*^*R239X*^ mice with an equal amount of 4-HB and β-RA incorporated into the chow. Interestingly, the co-administration of 4-HB and β-RA suppressed the mild inhibitory effect of β-RA over CoQ_9_ biosynthesis in the skeletal muscle (Fig. 4D) and CoQ_10_ biosynthesis in the brain, kidneys, and liver (Fig. 4F-H) of *Coq9*^*+/+*^ mice (compare with Fig. 3). Moreover, CoQ_9_ increased in the brain (Fig. 4A) and the kidneys (Fig. 4B) of *Coq9*^*+/+*^ mice treated with the combination of 4-HB and β-RA compared to the untreated *Coq9*^*+/+*^ mice. In *Coq9*^*R239X*^ mice, the untreated and treated groups showed similar levels of both CoQ_9_ (Fig. 4A-E) and CoQ_10_ (Fig. 4F-J) in all tissues. Importantly, the reduction of the levels of DMQ_9_ and of the DMQ_9_/CoQ_9_ ratio induced by β-RA (Fig. 3, S3 and S4) in *Coq9*^*R239X*^ mice seemed to be suppressed by the co-administration of 4-HB and β-RA (Fig. 4K-T). Consequently, the co-administration of 4-HB and β-RA suppressed the increase of survival of *Coq9*^*R239X*^ mice found after the treatment with β-RA alone (Fig. 4U). Together, these data demonstrate that β-RA acts therapeutically in *Coq9*^*R239X*^ mice by entering the CoQ biosynthetic pathway, leading to a reduction in the levels of DMQ_9_.

**Figure 4.**
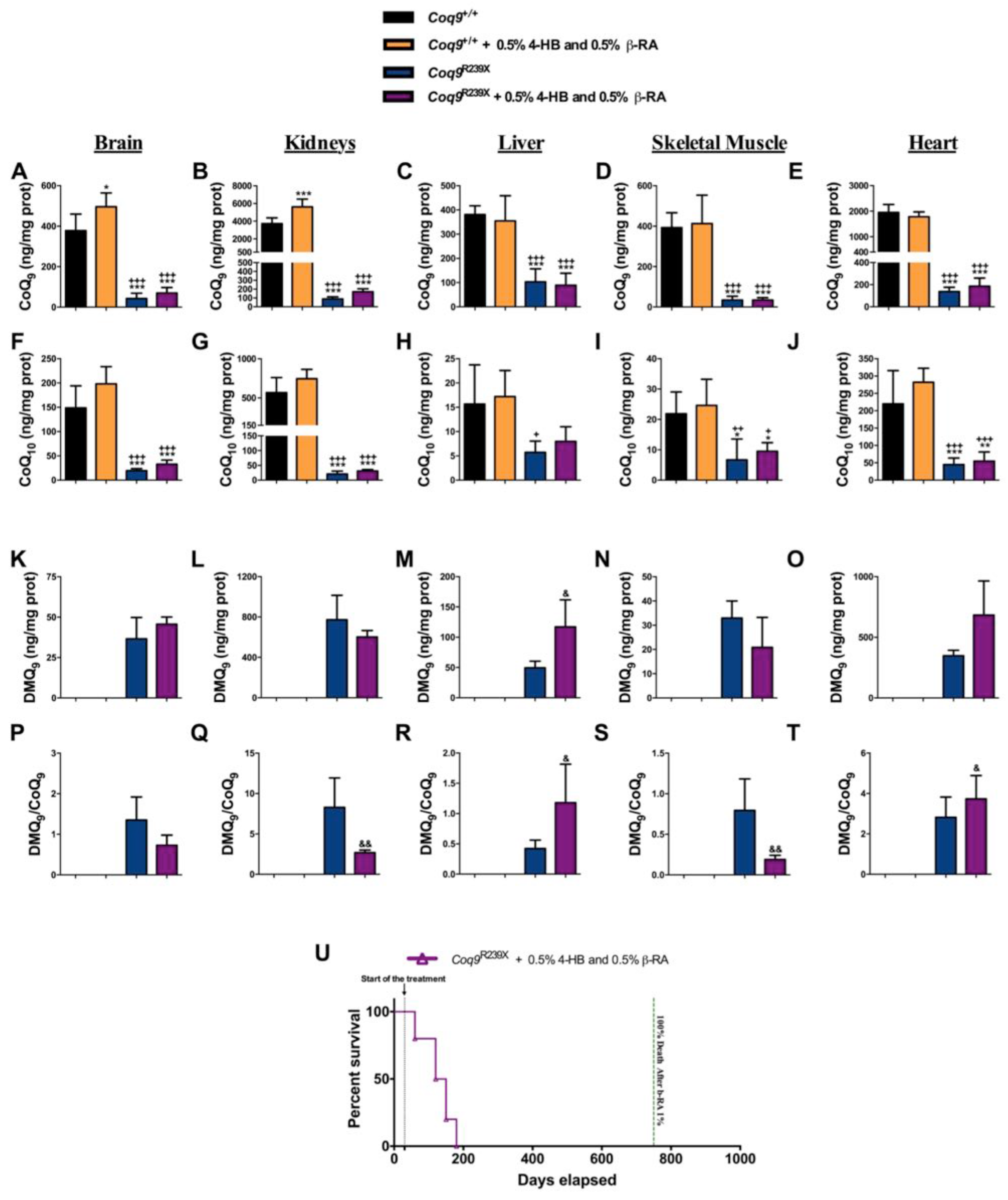
Co-administration of 4-HB suppresses the effects of β-RA treatment in *Coq9*^*+/+*^ and *Coq9*^*R239X*^ mice. (**A** to **E**) Levels of CoQ_9_ in the brain (A), kidneys (B), liver (C), skeletal muscle (D) and heart (E) from *Coq9*^*+/+*^ mice, *Coq9*^*+/+*^ mice under 0.5% 4-HB + 0.5% β-RA treatment, *Coq9*^*R239X*^ mice, and *Coq9*^*R239X*^ mice under 0.5% 4-HB + 0.5% β-RA treatment. (**F** to **J**) Levels of CoQ10 in the brain (F), kidneys (G), liver (H), skeletal muscle (I) and heart (J) from *Coq9*^*+/+*^ mice, *Coq9*^*+/+*^ mice under 0.5% 4-HB + 0.5% β-RA treatment, *Coq9*^*R239X*^ mice, and *Coq9*^*R239X*^ mice under 0.5% 4-HB + 0.5% β-RA treatment. (**K** to **O**) Levels of DMQ9 in the brain (K), kidneys (L), liver (M), skeletal muscle (N) and heart (O) from *Coq9*^*+/+*^ mice, *Coq9*^*+/+*^ mice under 0.5% 4-HB + 0.5% β-RA treatment, *Coq9*^*R239X*^ mice, and *Coq9*^*R239X*^ mice under 0.5% 4-HB + 0.5% β-RA treatment. (**P** to **T**) Ratio DMQ9/ CoQ9 in the brain (P), kidneys (Q), liver (R), skeletal muscle (S) and heart (T) from *Coq9*^*+/+*^ mice, *Coq9*^*+/+*^ mice under 0.5% 4-HB + 0.5% β-RA treatment, *Coq9*^*R239X*^ mice and *Coq9*^*R239X*^ mice, under 0.5% 4-HB + 0.5% β-RA treatment. (**U**) Survival curve of *Coq9*^*R239X*^ mice under 0.5% 4-HB + 0.5% β-RA treatment. Tissues from mice at 3 months of age. Data are expressed as mean ± SD. *P < 0.05; **P < 0.01; ***P < 0.001; differences versus *Coq9*^*+/+*^. +P < 0.05; ++P < 0.01; +++P < 0.001, differences *versus Coq9*^+/+^ after 0.5% 4-HB and 0.5% β-RA treatment. &P < 0.05; &&P < 0.01, differences *versus Coq9^R239X^* (one-way ANOVA with a Tukey’s post hoc test or t-test; n = 5–10 for each group).

### A metabolic switch in wild-type animals contributes to the effects of β-RA in reducing WAT

Since the interference of β-RA in CoQ metabolism in wild-type mice is very mild, the profound reduction of WAT is not likely attributed to CoQ metabolism. Thus, we investigated whether β-RA can target other mitochondrial pathways by performing quantitative proteomics on mitochondrial fractions of kidneys from wild-type mice treated with 1% β-RA for only two months and compare the results to those of kidneys from untreated wild-type mice (Data File S1). We chose a higher dose to ensure to the effects of the β-RA supplementation were clearly evident. Also, the analysis was done in the kidneys because this tissue maintained the highest levels of β-RA after the supplementation. In the kidneys of wild-type mice treated with β-RA compared to kidneys of untreated wild-type mice, 442 mitochondrial proteins were differentially expressed (Data File S2), with 300 proteins being over-expressed and 142 proteins being under-expressed. Canonical metabolic analysis showed enrichment (top 10) of pathways of fatty acid β oxidation, acetyl-CoA biosynthesis, the TCA cycle and the 2-ketoglutarate dehydrogenase complex, as well as enrichment of the related branched-chain α-keto acid dehydrogenase complex (Fig. 5A). Importantly, the prediction z-score revealed an inhibition of fatty acid β oxidation and an activation of acetyl-CoA biosynthesis and the TCA cycle (Fig. 5A), which is consistent with the changes found in the levels of key proteins in these pathways (Fig. 5B). Western blotting for the proteins ALDH1B1, GSK3β, EHHADH, and ACADM from mice fed both a 1% and 0.33% β-RA diet (Fig. 5C-D) validated these findings in the kidneys. Taken together, the results of the mitochondrial proteome analysis suggest that β-RA treatment stimulates the production and use of acetyl-CoA in the kidneys while repressing fatty acid β oxidation in the kidneys (Fig. 5E). Thus, we hypothesized that β-RA supplementation induces glycolysis at the expense of fatty acid β oxidation. For that, lipolysis may induce an increase of glycerol-3-P (G3P), which may stimulate glycolysis to provide the substrate for acetyl-CoA biosynthesis. Accordingly, the activities of the glycolytic enzymes phosphofructokinase (PFK) and pyruvate kinase (PK) were partially increased with the β-RA treatment (Fig. 5F-G). Also, G3P were increased with the β-RA treatment (Fig. 5H), while the levels of β-hydroxybutyrate (BHB) showed a notable but statistically insignificant increase with the β-RA treatment (Fig. 5I).

**Figure 5.**
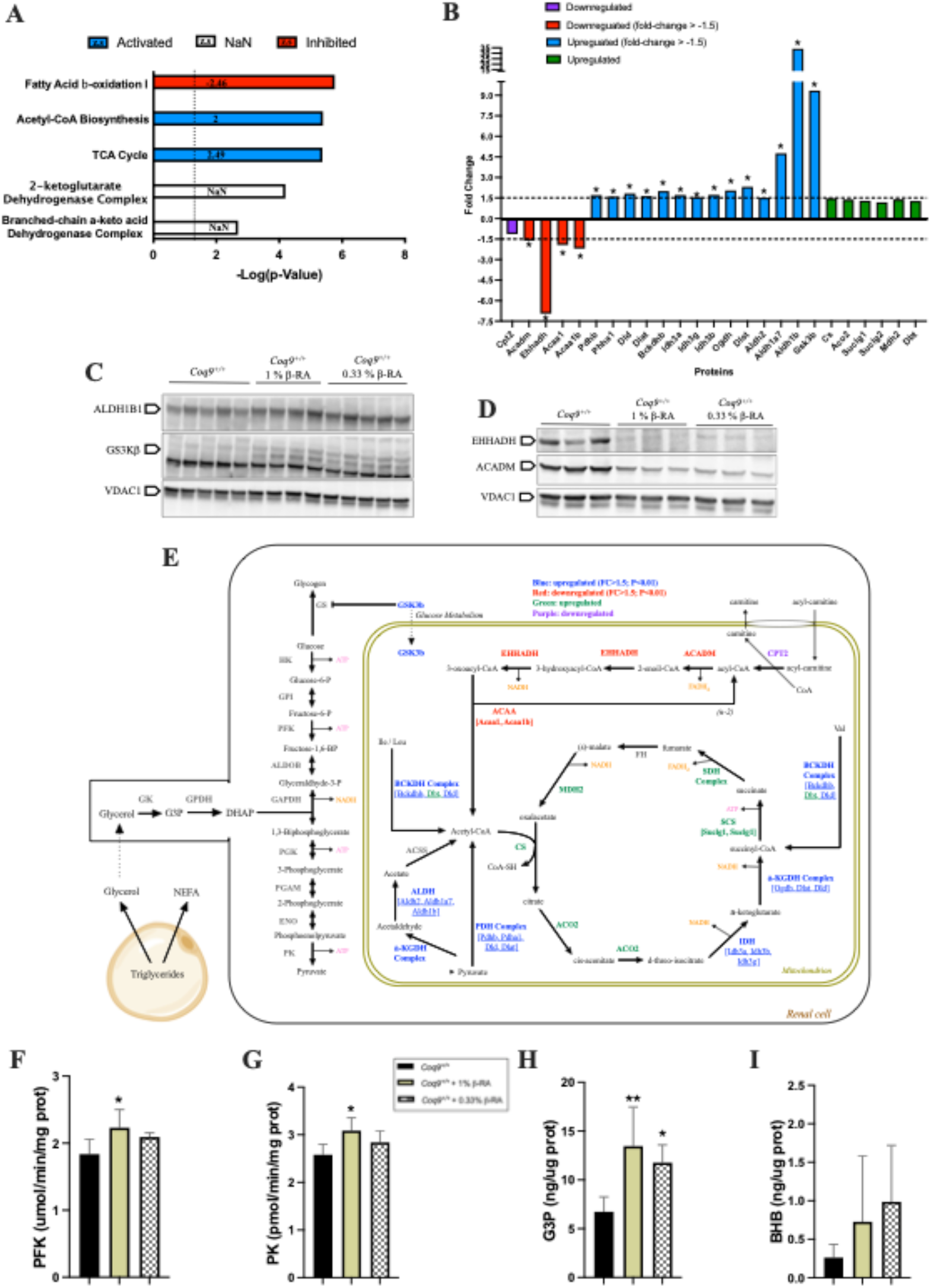
Adaptation of the mitochondrial proteome to the treatment with β-RA in the kidneys of *Coq9*^*+/+*^ mice. (**A**)Top enriched metabolic canonical pathways in the renal mitochondrial proteome from *Coq9*^*+/+*^ mice after two months of 1% β-RA supplementation. Dotted line: Adjusted *p* = 0.05. Blue signifies that the category is expected to be activated according to the z-score; red signifies that the category is expected to be inhibited according to the z-score. (**B**) Fold-change (treated/untreated) of the proteins involved in the identified enriched metabolic canonical pathways in the renal mitochondrial proteome. Purple signifies proteins that are downregulated; red signifies proteins that are downregulated with a fold-change > 1.5; blue signifies proteins that are upregulated with a fold-change > 1.5; green signifies proteins that are upregulated. *P<0.05. Mitochondrial proteomics was performed in isolated mitochondria. (**C** and **D**) Western-blot of some key proteins identified in the proteomics analysis to validate the changes observed with the treatment of β-RA. The validation is performed with the treatment of β-RA at 1% and extended to the treatment of β-RA at 0.33%. The selected proteins are ALDH1B1, GS3Kβ, EHHADH and ACADM. VDAC1 was used as loading control. The experiments were performed in tissue homogenate. (**E**) Schematic figure of the most important changes in the mitochondrial proteomes from the kidneys of *Coq9*^*+/+*^ mice after β-RA treatment. (**F** and **G**) Activities of the glycolytic enzymes Phosphofructokinase (PFK) (F) and Pyruvate Kinase (PK) (G) in the kidneys of *Coq9*^*+/+*^ mice treated with β-RA at 1% and 0.33%. (**H** and **Y**) Levels of Glycerol-3-Phosphate (G3P) (H) and β-hydroxybutyrate (BHP) (I) in the kidneys of *Coq9*^*+/+*^ mice treated with β-RA at 1% and 0.33%. Tissues from mice at 3 months of age. Data are expressed as mean ± SD. *P < 0.05; **P < 0.01, differences *versus Coq9*^*+/+*^(one-way ANOVA with a Tukey’s post hoc test or t-test; n = 5–7 for each group).

We performed similar analyses in the liver and skeletal muscle, two relevant tissues in the regulation of systemic energy metabolism, to check whether this metabolic switch was a common phenomenon. The levels of the proteins ALDH1B1, GSK3β, EHHADH, and ACADM in the liver and skeletal muscle did not change like the changes observed in the kidneys (Fig. 6A-F; Fig. S7A-F). However, PFK activity increased with the β-RA treatment in both tissues (Fig. 6G; Fig. S7G), suggesting an activation of glycolysis despite a lack of change of PK activity from the treatment (Fig. 6H; Fig. S7H). Also, G3P increased in the liver with the treatment of 1% β-RA, although these levels did not change at the low dose nor in the skeletal muscle with both doses (Fig. 6I, Fig. S7I). In the liver, the levels of BHB showed an observable but statistically insignificant increase with the β-RA treatment (Fig. 6J). The levels of *Fgf21*, a secretory endocrine factor that can affect systemic glucose and lipid metabolism (Salminen, Kaarniranta et al., 2017), trended upward with the β-RA treatment (Fig. 6K). The increase in BHB levels was also observed in the blood plasma with the treatment of 1% β-RA (Fig. 6L). However, the levels of non-esterified fatty acids (NEFA), which are products of lipolysis, were similar in the treated and untreated animals (Fig 6M). Also, the levels of glucagon, insulin, and the insulin/glucagon ratio were similar in the treated and untreated animals, which most likely reflects a homeostatic status with chronic administration of β-RA (Fig. 6N-P). These results suggest that metabolism in the kidneys and, to a lesser extent, the liver, contributes to the reduced WAT induced by β-RA in wild-type animals.

**Figure 6.**
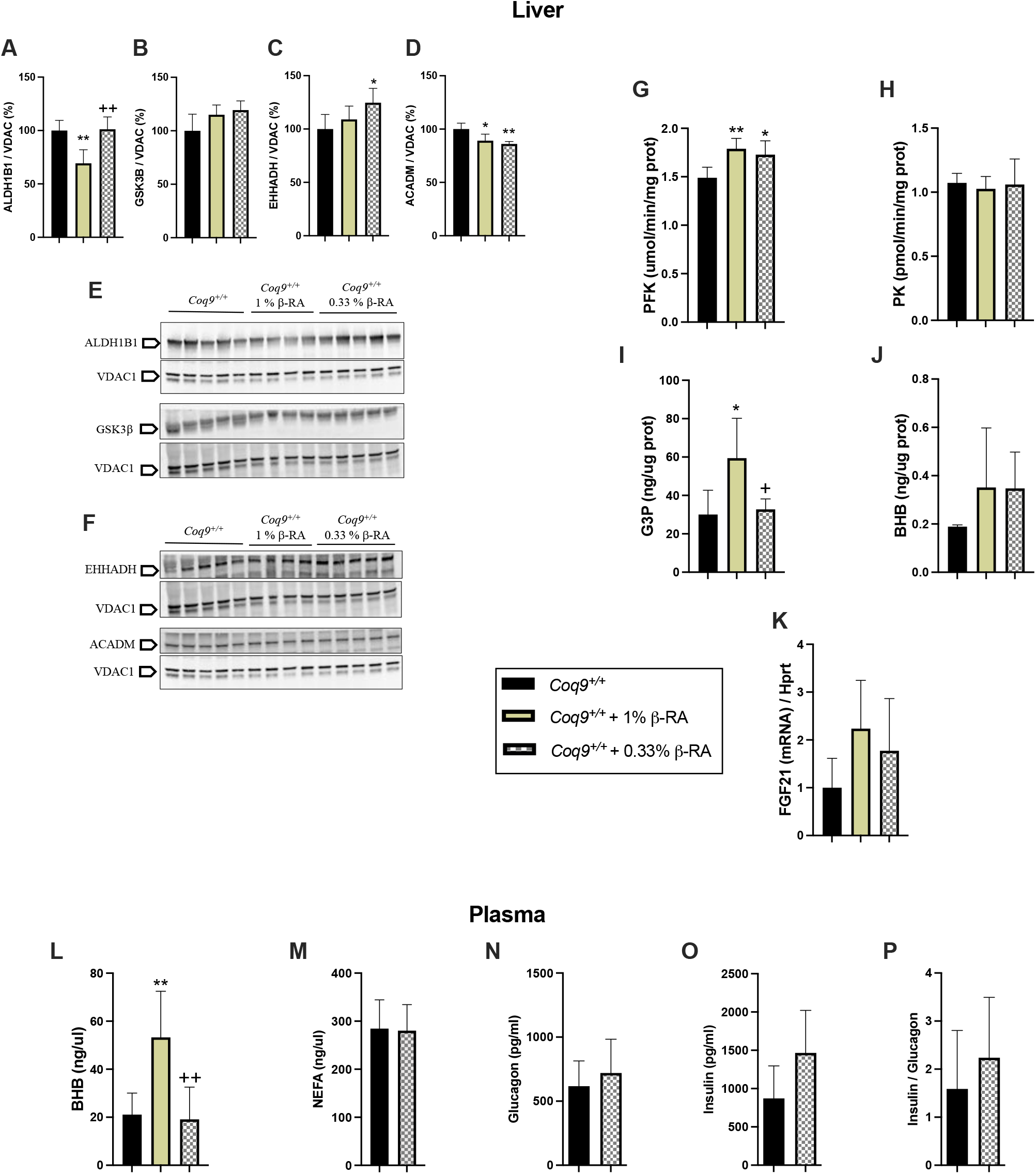
Metabolic characterization of liver and plasma after the treatment with β-RA in *Coq9*^*+/+*^ mice. (**A** to **F)** Levels of the proteins ALDH1B1 (A and E), GSK3β (B, and E), EHHADH (C and F) and ACADM (D and F) in the liver of *Coq9*^*+/+*^ mice treated with β-RA at 1% and 0.33%. VDAC1 was used as loading control. The experiments were performed in tissue homogenate. (**G** to **J**)Activities of the glycolytic enzymes Phosphofructokinase (PFK) (G) and Pyruvate Kinase (PK) (H) in the liver; levels of Glycerol-3-Phosphate (G3P) in the liver (I); levels of β-hydroxybutyrate (BHP) in the liver (J). (**K**) Levels of the FGF21 mRNA in the liver. (**L** to **P**) Levels of β-hydroxybutyrate (BHP) (L), non-esterified fatty acids (NEFA) (M), glucagon (N) and insulin (O) in the plasma of *Coq9*^*+/+*^ mice treated with β-RA; glucagon/insulin ratio (P) in the plasma of *Coq9*^*+/+*^ mice treated with β-RA. Tissues from mice at 3 months of age. Data are expressed as mean ± SD. *P < 0.05; **P < 0.01, differences *versus Coq9*^*+/+*^. +P < 0.05; ++P < 0.01 differences versus *Coq9*^*+/+*^ after 1% β-RA treatment (one-way ANOVA with a Tukey’s post hoc test or t-test; n = 5–7 for each group).

### β -RA directly inhibits adipogenesis

While the metabolic switch in the kidneys, and, to a lesser extent, the liver, may contribute to the utilization of energetic substrates that prevent the accumulation of WAT, we also wondered whether β-RA directly affects adipocytes. Thus, we treated 3T3-L1 preadipocytes with β-RA. In proliferative conditions, β-RA decreased cell proliferation (Fig. 7A,D), most likely due to an increase of p27 (Fig. 7B), a protein which inhibits the cell cycle progression at G1 (Cheng, Olivier et al., 1999, Ferguson, Nam et al., 2016). We also observed a decrease in CYCA2, a protein which promotes the division of the cells (Ferguson et al., 2016) (Fig. 7B). These changes in p27 and CYCA2 were not observed in differentiated 3T3-L1 cells (Fig. 7C) nor in C2C12 myoblasts under both proliferative and differentiative conditions (Fig. S8), indicating a cell-type specific effect. Consistently, 3T3-L1 cells treated with β-RA produced less fat (Fig. 7E-F), a phenomenon that may be mediated by the decrease in PPARγ levels (Figure 7G), and the upward trend of PPARδ levels (Figure 7H), two receptors that regulate adipogenesis (Lehrke & Lazar, 2005, Schoonjans, Staels et al., 1996). The decreased levels of CoQ_9_ (Fig. 7I-J) due to the competitive inhibition of CoQ biosynthesis induced by β-RA in control cells, a fact that has been previously reported (Luna-Sanchez et al., 2015, Pierrel, 2017), could also contribute to the decreased proliferation and fat production of 3T3-L1 cells (Carriere, Fernandez et al., 2003, Liu, Lin et al., 2012).

**Figure 7.**
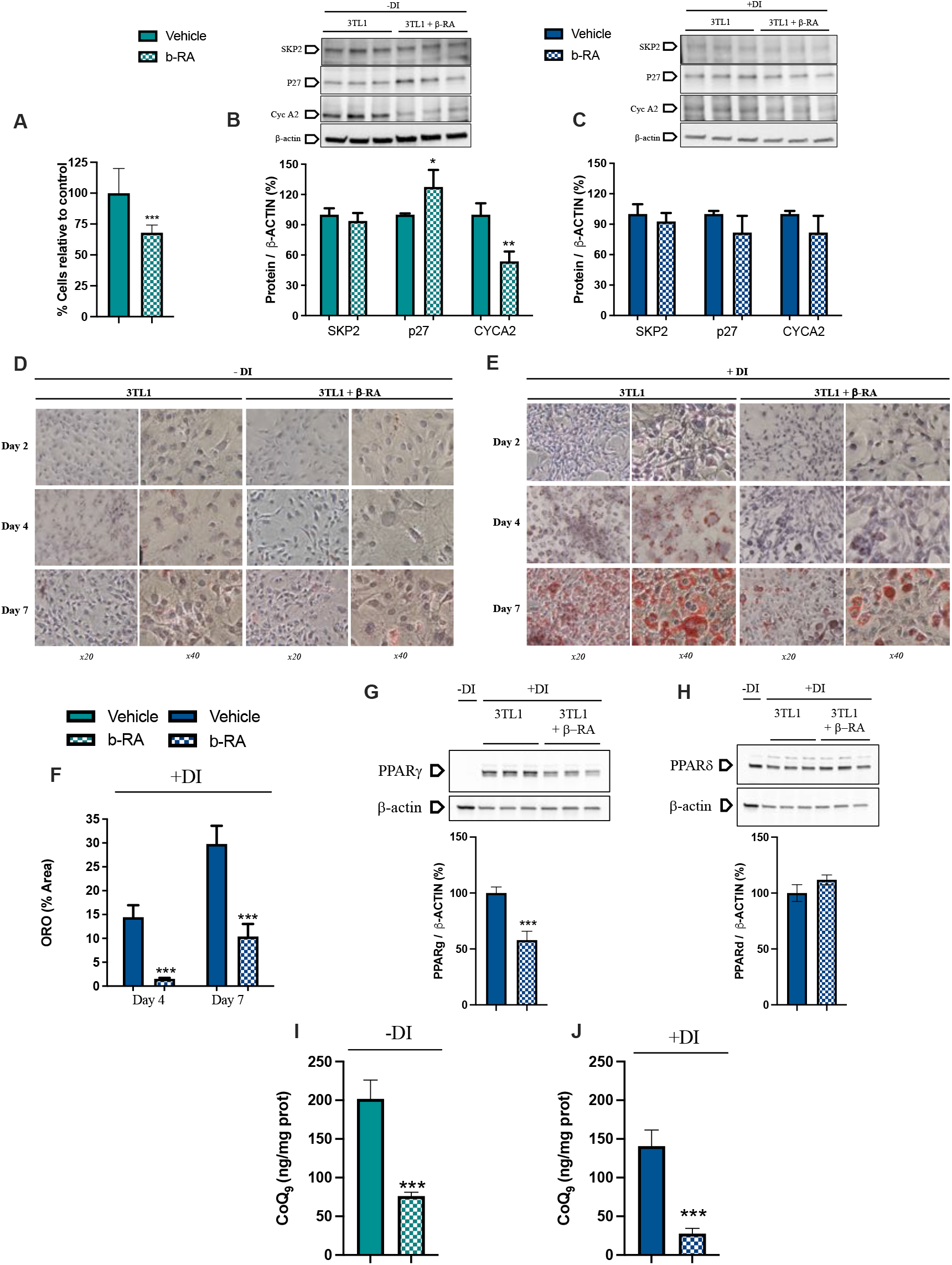
Direct effects of β;-RA on adipogenesis. (**A**) Percentage of 3TL1 cells after seven days of treatment with 1mM β;-RA, relative to the number of untreated 3TL1 cells. Cells cultured in proliferative conditions. (**B**) Levels of the proteins SKP2, p27 and CYCA2, which are involved in the control of the cell cycle. 3TL1 were treated for seven days with 1mM β;-RA in proliferative conditions. (**C**) Levels of the proteins SKP2, p27 and CYCA2, which are involved in cell cycle control. 3TL1 cells were treated for seven days with 1mM β;-RA in differentiative conditions. (**D** and **E**) Oil-red O staining in 3TL1 cells cultured under proliferative (D) and proliferative + differentiative (F) conditions. 3TL1 were treated with 1mM β;-RA from day 0 in both conditions and the stain was performed at three different days (2, 4 and 7). (**F**) Percentage of the area corresponding to the Oil Red O stains in 3TL1 cells in differentiative conditions after four and seven of treatment with 1mM β;-RA. (**G** and **H**) Levels of PPARγ and PPARδ in 3TL1 cells cultured proliferative + differentiative (F) conditions and treated with 1 mM β;-RA. The results in non-differentiated cells are shown in line one as negative control. (**I** and **J**) Levels of CoQ_9_ in 3TL1 cells cultured proliferative conditions (I) and differentiative conditions (J) and treated with 1 mM β;-RA. Data are expressed as mean ± SD. *P < 0.05; **P < 0.01; ***P < 0.001, differences *versus* untreated cells (t-test; n = 6 for each group).

Because other HBAs, e.g., salicylic acid or vanillic acid, can activate AMPK (Hawley, Fullerton et al., 2012, Jung, Park et al., 2017), an enzyme that plays a key role in cellular energy homeostasis (Garcia & Shaw, 2017, Herzig & Shaw, 2018), we investigated whether the observed effects of β-RA in WAT are due to the activation of AMPK through its phosphorylation. Thus, we quantified the levels of AMPK and p-AMPK, as well as two of its target proteins, ULK1/p-ULK1 and ACC/p-ACC, in WAT of wild-type mice at 18 months of age. Both the phosphorylated and unphosphorylated forms of the three proteins were increased, although the p-AMPK/AMPK, p-ULK1/ULK and p-ACC/ACC ratios were similar in untreated and treated animals (Fig. S9A-C), suggesting that AMPK is not a direct target of β-RA. Also, 3T3-L1 cells treated with β-RA did not experience changes in p-AMPK/AMPK ratio, with p-AMPK being almost undetectable in both treated and untreated cells (Fig. S9D-E).

## Discussion

β;-R -RA is an HBA that shows powerful therapeutic benefits in CoQ deficiency mouse models caused by mutations in *Coq6, Coq7, Coq8b*, or *Coq9* (Hidalgo-Gutierrez et al., 2019, Wang et al., 2015, Widmeier et al., 2019a, Widmeier et al., 2019b). Those studies administered high doses of oral β;-RA, but the mechanisms have not been clearly elucidated in podocyte-specific *Coq6* or *Coq8b* knockout mice (Widmeier et al., 2019a, Widmeier et al., 2019b). Moreover, chronic β;-RA supplementation maintains wild-type mice with a lower body weight than those untreated (Wang et al., 2015), but the causes and mechanisms of this effect were completely unknown. In our current work, we demonstrate that the therapeutic mechanism of β;-RA in *Coq9*^*R239X*^ mice is based on the capability of this molecule to enter the CoQ biosynthetic pathway and compete with 4-HB, resulting in a reduction of the levels of DMQ, an intermediate metabolite that is detrimental for mitochondrial function (Yang, Vasta et al., 2011). Moreover, our study reveals that β;-RA prevents the accumulation of WAT during the animal development and aging, thus preventing age-related hepatic steatosis. This powerful effect is due to an inhibition of preadipocyte proliferation and fat production, as well as a stimulation of lipolysis, gluconeogenesis, and glucose and acetyl-CoA utilization, mainly in the kidneys.

The fundamental rationale of the treatment with β;-RA in primary CoQ deficiency is the induction of a bypass effect, since β;-RA has the hydroxyl group that is normally incorporated into the benzoquinone ring by the hydroxylase COQ7. Because COQ9 is essential for the stability and function of COQ7 (Garcia-Corzo et al., 2013), defects in either *Coq7* or *Coq9* are susceptible to be effectively treated by β;-RA (Freyer, Stranneheim et al., 2015, Hidalgo-Gutierrez et al., 2019, Luna-Sanchez et al., 2015, Wang et al., 2015, Wang et al., 2017). Surprisingly, β;-RA treatment was also successful in podocyte-specific *Coq6* or *Coq8b* knockout mice, yet the mechanism in those cases were apparently not related to a bypass effect, suggesting the β;-RA may induce additional therapeutic mechanisms. Our results, however, confirm that the therapeutic mechanism of β;-RA in *Coq9*^*R239X*^ mice is due to its action in CoQ metabolism as demonstrated by: 1) the decrease in the levels of DMQ, with the effect more intense in the kidneys (the tissue that accumulated more β;-RA); and 2) the suppression of the therapeutic effect of β;-RA by the co-administration of 4-HB, which attenuated the decrease of DMQ_9_, thus supporting the theory of competition between the molecules to enter the CoQ biosynthetic pathway *in vivo* (Pierrel, 2017). The results obtained with the co-administration of 4HB and β;-RA also suggest that the K_M_ for β;-RA is higher than the K_M_ for 4-HB in the prenylation reaction catalyzed by COQ2 (Hidalgo-Gutierrez et al., 2019, Pierrel, 2017). Moreover, the therapeutic effects observed in this study were achieved with a third of the dose previously used (Hidalgo-Gutierrez et al., 2019). Thus, the effects in this study were also similar to the results published in the *Coq7* conditional KO mice (Wang et al., 2017) despite the phenotypes of both models being substantially different (Garcia-Corzo et al., 2013, Wang et al., 2015). This is important because animal studies that use lower doses of a drug could potentially be translatable to the human situation, decreasing the cost of the treatment and being more feasible its administration, especially in the pediatric population. Our results in the *Coq9*^*R239X*^ mice, however, show that β;-RA has limitations in inducing an increase in the levels of CoQ, suggesting that the co-supplementation of β;-RA and CoQ_10_ could result in improved therapeutic outcomes (Gonzalez-Garcia, Hidalgo-Gutierrez et al., 2020).

In wild-type animals, chronic β;-RA supplementation prevented the accumulation of WAT. The *in vitro* experiments in this study demonstrated that β;-RA inhibits preadipocytes proliferation, a result that has also been achieved by other phenolic acids (Aranaz, Navarro-Herrera et al., 2019, Hsu & Yen, 2007), including *p*-coumaric (Aranaz et al., 2019), which has been reported to serve as benzoquinone precursor for CoQ biosynthesis in humans and mice (Xie, Williams et al., 2015). Whether the alteration on CoQ biosynthesis induced by β;-RA, i.e. the decrease in CoQ levels or the mild accumulation of DMQ, may contribute to the accumulation of WAT remain to be elucidated. The anti-proliferative effect of β;-RA in preadipocytes induces the downregulation of PPARγ, which seems to be critical for the suppression of adipocyte differentiation and the development of mature adipocytes (Rosen & Spiegelman, 2000). Consequently, β;-RA may act by preventing WAT hyperplasia and hypertrophy, both of which contribute to avoid overweight and obesity in children and adults (Jo, Gavrilova et al., 2009, Landgraf, Rockstroh et al., 2015, Tchoukalova, Votruba et al., 2010).

In addition to the direct effects of β;-RA in adipocytes, *in vivo* experiments utilizing hypotheses generated by proteomic profiling, and following these observations up with focused validation experiments, showed a tissue metabolic switch, mainly in the kidneys. This tissue could account for up to 40% of the overall gluconeogenesis of the body under certain conditions, e.g. the post-absorptive phase (Gerich, Meyer et al., 2001, Legouis, Faivre et al., 2020), during which glycerol is one of the gluconeogenic renal precursors (Gerich et al., 2001). Although renal gluconeogenesis mainly serves to produce glucose only for its own utilization in the kidneys, this metabolic process can also participate in the regulation of systemic glucose metabolism (Legouis et al., 2020). Therefore, our results suggest that the β;-RA induces renal gluconeogenesis from glycerol, and the resulting glucose is used in glycolysis to produce pyruvate and then acetyl-CoA, which is ultimately funneled into the TCA cycle. Acetyl-CoA may not only be produced through the classical pathway but also through an alternative pathway that involves α-ketoglutarate dehydrogenase and aldehyde dehydrogenase and uses acetaldehyde as intermediate metabolite (Liu, Cooper et al., 2018b). Interestingly, the production and use of acetyl-CoA in mitochondria has been postulated as a metabolic signal of survival in the organism (Shi & Tu, 2015), which is consistent with a reduction in the WAT content (Huang, Zhang et al., 2018, Shi & Tu, 2015), the stimulation of ketogenesis (Fletcher, Deja et al., 2019, Shi & Tu, 2015), the limitation of fatty acid synthesis, and the prevention of hepatic steatosis (Fletcher et al., 2019, Huang et al., 2018, Shi & Tu, 2015). Nevertheless, it is unclear whether the metabolic effects in the kidneys, and to a lesser extent, in the liver, are due to β;-RA itself or if they are the consequences of having a low amount of WAT. This second option could explain the downregulation of fatty acid β;-oxidation in the kidneys and the subsequent preference for glucose metabolism. A potential regulator for all these metabolic changes is GSK3β;, which is highly increased in the mitochondria of the treated wild-type animals. GSK3β; regulates a variety of cellular processes, including glucose metabolism. In fact, its upregulation has been associated with an amelioration of the diabetes-induced kidney injury (Mariappan, Prasad et al., 2014). Consequently, these metabolic adaptations in the kidneys in response to chronic supplementation of β;-RA could explain, at least in part, the positive therapeutic outcomes achieved in the podocyte-specific *Coq6* or *Coq8b* knockout mice (Widmeier et al., 2019a, Widmeier et al., 2019b) and open the potential application of β;-RA in treating other renal metabolic diseases.

To conclude, the results reported here demonstrate that chronic supplementation with β;-RA in mice induces different metabolic effects with relevant therapeutic implications for the treatment of primary CoQ deficiency and the prevention of age-related overweight and associated hepatic steatosis. The first application is based on the ability of β;-RA to enter the CoQ biosynthetic pathway, compete with a lower affinity with the natural substrate 4-HB and, consequently, reduce the levels of DMQ in cases of defects in *Coq9* or *Coq7*. The second application is based on a combination of direct influences over WAT, ultimately preventing the hyperplasia and hypertrophy of adipocytes, and to indirect systemic mechanisms, mainly by adaptations of renal metabolism. Nevertheless, this study has some limitations: 1) although β;-RA can prevent the accumulation of WAT during aging, it is unknown whether it can reduce WAT in already obese animals; 2) although this long-term study shows convincing therapeutic actions of β;-RA, the effects of β;-RA administration should be evaluated in mice with different genetic backgrounds and models of both diet-induced obesity and genetic-induced obesity; and 3) a minimal effective dose and potential dose-dependent specific effects must be defined for both therapeutic applications. Nevertheless, the data of the present work are relevant for the future translation of the treatment with β;-RA into the clinic, especially considering that we have shown the effects of the long-term administration of β;-RA in a mouse model of age-related overweight and mitochondrial encephalopathy due to CoQ deficiency.

## Materials and Methods

### Animals and treatments

*Coq9*^+/+^ and *Coq9* ^*R239X*^ mice were used in the study, both of which harboured a mix of C57BL/6N and C57BL/6J genetic background. The *Coq9*^*R239X*^ mouse model (MGI: 5473628) was previously generated and characterized (Garcia-Corzo et al., 2013, Luna-Sanchez et al., 2015, Luna-Sanchez et al., 2017). All animal manipulations were performed according to a protocol approved by the Institutional Animal Care and Use Committee of the University of Granada (procedures numbers 18/02/2019/016 and 16/09/2019/153) and were in accordance with the European Convention for the Protection of Vertebrate Animals used for Experimental and Other Scientific Purposes (CETS #123) and the Spanish law (R.D. 53/2013). Mice were housed in the Animal Facility of the University of Granada under an SPF zone with lights on at 7:00 AM and off at 7:00 PM. Mice had unlimited access to water and rodent chow (SAFE® 150, which provides 21%, 12.6% and 66.4% energy from proteins, lipids and nitrogen-free extracts, respectively). Unless stated otherwise, the analytical experiments were completed in animals at 3 or 18 months of age.

β;-Resorcylic acid (β;-RA) was given to the mice in the chow at a concentration of 0.33 % (w/w). For some experiments, a concentration of 1% (w/w) β;-RA was used for two months (Hidalgo-Gutierrez et al., 2019). A mix of β;-RA and 4-HB (at a concentration of 0.5% each one) was also provided in the chow for particular experiments. Mice began receiving the assigned treatments at 1 month of age, and the analyses were performed at the age indicated for each case. Animals were randomly assigned to experimental groups. Data were randomly collected and processed.

The body weights were recorded once a month. To weight skeletal muscle, mice were sacrificed at 18 month of age and the *gastrocnemius* and *vastus lateralis* were dissected and weighed in a laboratory scale. To weigh WAT, mice were sacrificed at 18 month of age and the epididymal, mesenteric and inguinal WAT were dissected and weighted on a laboratory scale.

The motor coordination was assessed at different months of age using the rotarod test by recording the length of time that mice could remain on the rod (“latency to fall”), rotating at a rate of 0.1 rpm/s for up to 300 s. Muscle strength was assessed using a computerized grip strength meter (Model 47200, Ugo-Basile, Varese, Italy). The experimenter held the mouse gently by the base of the tail, allowing the animal to grab the metal bar with the forelimbs before being gently pulled until it released its grip. The peak force of each measurement was automatically recorded by the device and expressed in grams (g). The hindlimb grip strength of each mouse was measured in duplicate with at least 1 min between measurements (Luna-Sanchez et al., 2015).

### Cell culture and cell assays

3T3-L1 preadipocytes (ECACC #: 86052701; lot CB 2618) were obtained from the cell bank of the University of Granada and maintained in DMEM containing 10% Fetal Calf Serum (FCS) in a humidified atmosphere of 5% CO_2_ at 37°C. The differentiation of the preadipocytes was induced 2 days post-confluence (day 0) following the manufacturer instructions (Sigma, DIF001-1KT) by the addition 0.5 mM IBMX, 1 μM dexamethasone and 10 μg/ml insulin [multiple daily insulin (MDI)] for 2 days. Subsequently, the culture medium was changed to DMEM and 10% Fetal Bovine Serum (FBS) containing insulin. After 2 days, the medium was replaced with DMEM and 10% FBS, and the cells were incubated for a further 2 days until the cells were harvested to be used in the experiments described below.

C2C12 myocytes (ECACC #: 91031101; lot 08F021) were obtained from the cell bank from the University of Granada and maintained in DMEM containing 10% FBS in a humidified atmosphere of 5% CO_2_ at 37°C. The differentiation of the preadipocytes was induced 1-day post-confluence (day 0) by the change to a 1% FBS medium. Subsequently, the culture medium was changed to DMEM and 1% FBS. The medium was changed every other day and the cells were harvested to be used in the experiments described below.

In both cell types, 3T3-L1 and C2C12, the assays were carried out in three experimental conditions: proliferative, differentiative or proliferative + differentiative. Proliferative conditions were developed in both type of cells after cell splitting, and cells were collected upon reaching the confluency at day 7. Differentiative conditions were initiated, in both cell types, when the cells reached confluency. In 3T3-L1 cells differentiation was induced with the differentiation medium described above. In C2C12, differentiation was induced in a medium with 1% FBS, as described above. Cells were collected at day 7. Proliferative + differentiative conditions combined both procedures in the same experiment. β;-RA was added at a final concentration of 1mM every other day in each experimental condition.

To visualize the lipid droplets, 3T3-L1cells were fixed in formalin and stained with oil red solution at days 2, 4 and 6, in both proliferative and proliferative + differentiative conditions. Cell viability and proliferation were quantified at day 7 using a Vybrant MTT Cell Proliferation Assay Kit according to the manufacturer instructions. Absorbance was measured at 450 nm on a microplate reader (Powerwave x340 spectrophotometer, Biotek).

### Histology and immunohistochemistry

Tissues were fixed in formalin and paraffin embedded. Multiple sections (4 μm thickness) were deparaffinized with xylene and stained with hematoxylin and eosin (H&E), Masson’s trichrome or oil red. Immunohistochemistry was carried out on the same sections, using the following primary antibodies: glial fibrillary acidic protein or anti-GFAP (Millipore, MAB360). Dako Animal Research Kit for mouse primary antibodies (Dako Diagnóstico S.A., Spain) was used for the qualitative identification of antigens by light microscopy. Sections were examined at 40–400 magnifications with a Nikon Eclipse Ni-U microscope, and the images were scanned under equal light conditions with the NIS-Elements Br computer software.

### Plasma and urine analysis

Blood samples were collected in K_3_-EDTA tubes using a golden rod lancet and the submandibular vein of mice as puncture site. The plasma was extracted from blood samples by centrifugation at 4,500g for 10 minutes at 4ºC. Biochemical analysis from urine and plasma were developed in a biochemical analyzer Bs-200 (Shenzhen Mindray Bio-Medical Electronics Co., Ttd) and reagents from Spinreact.

The NEFAS concentration was quantified using the Free Fatty Acid Quantitation Kit (MAK044) according to the technical bulletin (Sigma-Aldrich, United States). The results are expressed in ng/μl.

The insulin concentration was quantified using the Mouse INS ELISA Kit (EM0260) according to the manufacturer’s instructions (FineTest, China). The results are expressed in pg/ml.

The Glucagon concentration was quantified using the Mouse GC ELISA Kit (EM0562) according to the manufacturer’s instructions (FineTest, China). The results are expressed in pg/ml.

### Mitochondrial proteomics analysis

Both *Coq9*^+/+^ mice and *Coq9* ^+/+^ mice under 1% of β;-RA supplementation were sacrificed, and the kidneys were removed and washed in saline buffer. The tissues were chopped with scissors in 3 ml HEENK (10 mM HEPES, 1 mM EDTA, 1 mM EGTA, 10 mM NaCl, 150 mM KCl, pH 7.1, 300 mOsm/l) containing 1 mM phenylmethanesulfonyl fluoride (from 0.1 M stock in isopropanol) and 1x protease inhibitor cocktail (Pierce). The tissues were homogenized with a 3 ml dounce homogenizer (5 passes of a tight-fitting Teflon piston). Each homogenate obtained was rapidly subjected to standard differential centrifugation methods until the mitochondrial pellet was obtained as previously described (Liu, Lossl et al., 2018a). Briefly, the homogenate was centrifuged at 600 *g* for 5 min at 4 °C (twice), and the resultant supernatant was centrifuged at 9,000 *g* for 5 min at 4 °C. The final pellet, corresponding to a crude mitochondrial fraction, was resuspended in 500 μl of HEENK medium without PMSF or protease inhibitor (Liu et al., 2018a). Protein concentration was determined (using Bradford dye (BIO-RAD) and a Shimadzu spectrophotometer, resulting in approximately 3 mg protein for renal mitochondria and 1.5mg for cerebral mitochondria. To verify the content of the mitochondrial fraction, Complex IV activity was determined by optical absorption of the difference spectrum at 550 nm, as previously described (Luna-Sanchez et al., 2017).

The purified mitochondria were spun down to remove the previous buffer, and lysis buffer (1% sodium deoxycholate SDC in 100 mM tris at pH 8.5) was added to the pellets. Samples were boiled for 5 minutes at 99ºC to denature all the proteins and then sonicated by microtip probe sonication (Hielscher UP100H Lab Homogenizer) for 2 min with pulses of 1s on and 1s off at 80% amplitude. Protein concentration was estimated by BCA assay and 200 µg were taken of each sample. 10 mM tris (2-carboxyethyl) phosphine and 40 mM chloroacetamide (final concentration) at 56 ºC were added to each of these 200 µg samples for 10 minutes to reduce and alkylate disulfide bridges. After this step, samples were digested with LysC (Wako) in an enzyme/protein ratio of 1:100 (w/w) for 1 h, followed by a trypsin digest (Promega) 1:50 (w/w) overnight. Protease activity was quenched with trifluoroacetic acid (TFA) to a final pH of ∼2. Samples were then centrifuged at 5,000*g* for 10 minutes to eliminate the insoluble SDC, and loaded on an OASIS HLB (Waters) 96-well plate. Samples were washed with 0.1% TFA, eluted with a 50/50 ACN and 0.1% TFA, dried by SpeedVac (Eppendorf, Germany), and resuspended in 2% formic acid prior to MS analysis.From each sample, 5 µg were taken from each sample and pooled in order to be used for quality control for MS (1 QC was analyzed every 12 samples) and to be fractionated at high-pH for the Match between runs.

All samples with the QC and 7 high-pH fractions were acquired using an UHPLC 1290 system (Agilent Technologies; Santa Clara, USA) coupled on-line to an Q Exactive HF mass spectrometer (Thermo Scientific; Bremen, Germany). Peptides were first trapped (Dr. Maisch Reprosil C18, 3 μm, 2 cm × 100 μm) prior to separation on an analytical column (Agilent Poroshell EC-C18, 2.7 μm, 50 cm × 75 μm). Trapping was performed for 5 min in solvent A (0.1% v/v formic acid in water), and the gradient was as follows: 10% – 40% solvent B (0.1% v/v formic acid in 80% v/v ACN) over 95 min, 40– 100% B over 2 min, then the column was cleaned for 4 min and equilibrated for 10 min (flow was passively split to approximately 300 nL/min). The mass spectrometer was operated in a data-dependent mode. Full-scan MS spectra from m/z 300-1600 Th were acquired in the Orbitrap at a resolution of 120,000 after accumulation to a target value of 3E6 with a maximum injection time of 120 ms. The 15 most abundant ions were fragmented with a dynamic exclusion of 24 sec. HCD fragmentation spectra (MS/MS) were acquired in the Orbitrap at a resolution of 30,000 after accumulation to a target value of 1E5 with an isolation window of 1.4 Th and maximum injection time 54 ms.

All raw files were analyzed by MaxQuant v1.6.10 software (Cox & Mann, 2008) using the integrated Andromeda Search engine and searched against the mouse UniProt Reference Proteome (November 2019 release with 55412 protein sequences) with common contaminants. Trypsin was specified as the enzyme, allowing up to two missed cleavages. Carbamidomethylation of cysteine was specified as fixed modification and protein N-terminal acetylation, oxidation of methionine, and deamidation of asparagine were considered variable modifications. We used all the automatic settings and activated the “Match between runs” (time window of 0.7 min and alignment time window of 20 min) and LFQ with standard parameters. The files generated by MaxQuant were opened from Perseus for the preliminary data analysis: the LFQ data were first transformed in log2, then identifications present in at least N (3/5) biological replicates were kept for further analysis; missing values were then imputed using the standard settings of Perseus. Ingenuity Pathway Analysis (IPA) analysis was used to identified changes in metabolic canonical pathways and their z-score predictions (Raimundo, Vanharanta et al., 2009).

### Sample preparation and western blot analysis in tissues and cells

For western blot analyses, a glass-Teflon homogenizer was used to homogenize mouse kidney, liver, skeletal muscle, and WAT samples at 1100 rpm in T-PER® buffer (Thermo Scientific) with protease and phosphatase inhibitor cocktail (Pierce). Homogenates were sonicated and centrifuged at 1000g for 5 min at 4 ºC, and the resultant supernatants were used for western blot analysis. For western blot analyses in cells, the pellets containing the cells were re-suspended in RIPA buffer with protease inhibitor cocktail. About 30 μg of protein from the sample extracts were electrophoresed in 12% Mini-PROTEAN TGXTM precast gels (Bio-Rad) using the electrophoresis system mini-PROTEAN Tetra Cell (Bio-Rad). Proteins were transferred onto PVDF 0.45-μm membranes using a Trans-blot Cell (Bio-Rad) and probed with target antibodies. Protein–antibody interactions were detected using peroxidase-conjugated horse anti-mouse, anti-rabbit, or anti-goat IgG antibodies and Amersham ECLTM Prime Western Blotting Detection Reagent (GE Healthcare, Buckinghamshire, UK). Band quantification was carried out using an Image Station 2000R (Kodak, Spain) and a Kodak 1D 3.6 software. Protein band intensity was normalized to VDAC1 for mitochondrial proteins and to GAPDH or β;-actin to for cytosolic proteins. The data were expressed in terms of percent relative to wild-type mice or control cells. The following primary antibodies were used: anti-ALDH1B1 (Proteintech, 15560-1-AP), anti-GSK3B (Proteintech, 22104-1-AP), anti-EHHADH (Santa Cruz, sc-393123), anti-ACADM (Abcam, ab110296), anti-SKP2 (Proteintech, 15010-AP), anti-P27 (Proteintech, 25614-1-AP), anti-Cyc A2 (Proteintech, 18202-1-AP), anti-β;-ACTIN (Santa Cruz, sc-47778), anti-PPARγ (Thermo Scientific, MA5-14889), anti-PPARδ (Thermo Scientific, PA1-823A), anti-AMPK (Cell Signaling, #2532), anti-P-AMPK (Cell Signaling, #2531), anti-ULK1 (Cell Signaling, #8054) anti-P-ULK1 (Cell Signaling, #5869) anti-ACC (Cell Signaling, #3676) anti-P-ACC (Cell Signaling, #11818).

### Quantification of CoQ_9_ and CoQ_10_ levels in mice tissues and 3T3-L1 cells

After lipid extraction from homogenized tissues and cultured cells, CoQ_9_ and CoQ_10_ levels were determined via reversed-phase HPLC coupled to electrochemical detection, as previously described (Garcia-Corzo et al., 2013, Luna-Sanchez et al., 2015). The results were expressed in ng CoQ/mg protein.

### CoQ-dependent respiratory chain activities

Coenzyme Q-dependent respiratory chain activities were measured in tissue samples of brain, kidney, skeletal muscle, and heart. Tissue samples were homogenized in CPT medium (0.05 M Tris-HCl, 0.15 M KCl, pH 7.5) at 1,100 rpm in a glass–Teflon homogenizer. Homogenates were sonicated and centrifuged at 600 g for 20 min at 4 °C, and the supernatants obtained were used to measure CoQ-dependent respiratory chain activities (CI + III and CII + III) as previously described (Hidalgo-Gutierrez et al., 2019).

### Metabolic assays in tissues

Phosphofructokinase enzyme activity was measured using kit from Sigma-Aldrich (Phosphofructokinase Activity Colorimetric Assay Kit MAK093) according to manufacturer instructions. The enzyme activity was expressed in μmol/min/mg protein.

Pyruvate kinase enzyme activity was measured using kit from Sigma-Aldrich (Pyruvate kinase Activity Colorimetric Assay Kit MAK072) according to manufacturer instructions. The enzyme activity was expressed in pmol/min/mg protein.

The G3P concentration was quantified using the Glycerol-3-Phosphate Assay Kit (MAK207) according to the technical bulletin (Sigma-Aldrich, United States). The concentration of G3P was expressed in ng/μg protein.

The BHB concentration was quantified using the Beta-Hydroxybutyrate Assay Kit (MAK041) according to the technical bulletin (Sigma-Aldrich, United States). The concentration of BHB was expressed in ng/μg protein.

### Mitochondrial respiration

Mitochondrial isolation from the brain and the kidneys was performed as previously described (Barriocanal-Casado, Hidalgo-Gutierrez et al., 2019, Hidalgo-Gutierrez et al., 2019). Mitochondrial respiration was measured with an XF^e24^ Extracellular Flux Analyzer (Seahorse Bioscience) (Barriocanal-Casado et al., 2019, Hidalgo-Gutierrez et al., 2019, Rogers, Brand et al., 2011). Mitochondria were first diluted in cold MAS 1X for plating (3.5 μg/ in brain; 2 μg/well in kidney). Next, 50 μl of mitochondrial suspension was delivered to each well (except for background correction wells) while the plate was on ice. The plate was then centrifuged at 2,000*g* for 10 min at 4°C. After centrifugation, 450 μl of MAS 1X + substrate (10 mM succinate, 2 mM malate, 2 mM glutamate and 10 mM pyruvate) was added to each well. Respiration by the mitochondria was sequentially measured in a coupled state with the substrate present (basal respiration or State 2) followed by State 3o (phosphorylating respiration, in the presence of ADP and substrates). State 4 (non-phosphorylating or resting respiration) was measured after addition of oligomycin when all ADP was consumed, and then maximal uncoupler-stimulated respiration was measured by FCCP (State 3u). Injections were as follows: port A, 50 μl of 40 mM ADP (4 mM final); port B, 55 μl of 30 μg/ml oligomycin (3 μg/ml final); port C, 60 μl of 40 μM FCCP (4 μM final); and port D, 65 μl of 40 μM antimycin A (4 μM final). All data were expressed in pmol/min/mg protein.

### Quantification of β;-RA and 4-HB levels in mice tissues

Tissues from mice were homogenized in water. The homogenate samples were then treated with a solution of methanol/water (80:20, v/v), shook for 1 minute, sonicated for 15 minutes and then centrifuged at 5,000*g* for 25 minutes at 4ºC (Borges et al, 2017).

The supernatants were analyzed using a Thermo Scientific™ UltiMate™ 3000 UHPLC system (Waltham, Massachusetts, United States), consisting of an UltiMate™ 3000 UHPLC RS binary pump and an UltiMate™ 3000 UHPLC sample manager coupled to a Thermo Scientific™ Q Exactive™ Focus Hybrid Quadrupole-Orbitrap™ detector of mass spectrometer (MS/MS) with an electrospray ionization in negative mode (Waltham, Massachusetts, United States). The analytical separation column was a Hypersil GOLD™ C18, 3 μm, 4.6 × 150 mm column (Thermo Scientific™) and the flow rate was 0.6 ml/min. The mobile phase consisted of two solutions: eluent A (H_2_O + 0.1% Formic acid, MS grade, Thermo Scientific™) and eluent B (acetonitrile + 0.1% Formic acid, MS grade, Thermo Scientific™). Samples were eluted over 30 min with a gradient as follow: 0 min, 95% eluent A; 0-25 min, 70% eluent A; 25-25.1 min, 95 % eluent A; 25.1-30 min, 95% eluent A. Capillary and auxiliary gas temperatures were set at 275 and 450 °C, respectively. Sheath gas flow rate used was at 55 arbitrary units, auxiliary gas flow rate used was at 15 arbitrary units, sweep gas flow was used at 3 arbitrary units. Mass spectrometry analyses were carried out in full scan mode between 110 and 190 uma. To quantify the levels of 4-HB and β;-RA, we used a standard curve with both compounds at a concentration of 100 ng/ml, 10 ng/ml and 1 ng/ml.

### Statistical analysis

Number of animals in each group were calculated in order to detect gross ∼60% changes in the biomarker measurements (based upon alpha=0.05 and power of beta=0.8). We used the application available at http://www.biomath.info/power/index.htm. Animals were genotyped and randomly assigned to experimental groups in separate cages by the technician of the animal facility. Most statistical analyses were performed using the Prism 9 scientific software. Data are expressed as the mean ± SD of five-ten experiments per group. A one-way ANOVA with a Tukey’s post hoc test was used to compare the differences between three experimental groups. Studies with two experimental groups were evaluated using unpaired Student’s t-test. A P value of < 0.05 was considered to be statistically significant. Survival curve was analyzed by log-rank (Mantel-Cox) and the Gehan-Breslow-Wilcoxon tests. The statistical tests used for the transcriptomics and proteomics analyses are described in their respective sections.

## Data availability

The mass spectrometry proteomics data have been deposited to the ProteomeXchange (http://www.proteomexchange.org/). Consortium via the PRIDE partner repository with the dataset identifier PXD018311.

## Acknowledgements

We thank Seth Joel Drey for the English editing. We are grateful to Ana Fernandez (Universidad de Granada) for her technical support at the facilities of bioanalysis. We thank members of the Heck Lab for their support in analyzing the proteomics samples. This work was supported by grants from Ministerio de Ciencia e Innovación, Spain, and the ERDF (grant number RTI2018-093503-B-100), from the Muscular Dystrophy Association (MDA-602322), from the Junta de Andalucía (grant number P20_00134), from the University of Granada (grant reference “UNETE”, UCE-PP2017-06) and by EPIC-XS, project number 823839, funded by the Horizon 2020 programme of the European Union. P.G.-G. is a “FPU fellow” from the Ministerio de Universidades, Spain. M.E.D.-C. is supported by the Muscular Dystrophy Association. E.B.-C. is supported by the Junta de Andalucía. A.H.-G. was partially supported by the “FPU program” and the research program from the University of Granada.

## Author Contributions

A.H.-G. led the study, developed the phenotypic and survival assay and the body weight measurements, conducted the tests to assess the mitochondrial bioenergetics, western blot analyses, enzymatic assays, cell culture experiments, UHPLC EC and MS analysis, IPA analyses, analyzed the results, designed the figures, and wrote the manuscript. E.B.-C. contributed to the mitochondrial assays, western blot analyses, qPCR analyses, enzymatic assays, the management of the mouse colony, the rotarod test, phenotyping, cell culture experiments, and critically reviewed the manuscript. M.E.D.-C. performed the morphological analyses and critically reviewed the manuscript. P.G.-G. contributed to the mitochondrial assays, proteomics experiments, management of the mouse colony, and critically reviewed the manuscript. R.Z. supervised the proteomics experiments and analyses, and critically reviewed the manuscript. D.A.-C. contributed to the discussion. L.C.L. conceived the idea for the project, supervised the experiments, and edited the manuscript. All authors critically reviewed the manuscript. The results shown in this article constituted a section of the A.H.-G. doctoral thesis at the University of Granada.

