## Supplemental Material for "β-RA targets mitochondrial metabolism and adipogenesis, leading to therapeutic benefits against CoQ deficiency and age-related overweight"

### Tables

**Table S1. Markers of hepatic and renal function in the plasma and urine from *Coq9*$^{+/+}$ mice and *Coq9*$^{+/+}$ mice under 0.33% of β-RA supplementation.** GOT = glutamate-oxaloacetate transaminase; GPT = glutamate-pyruvate transaminase; AP = alkaline phosphatase.

| **PLASMA** | | |
|---|---|---|
| | ***Coq9*$^{+/+}$** | ***Coq9*$^{+/+}$ + 0.33 % β-RA** |
| | Mean ± SD | |
| GOT (U/L) | 76.20 ± 21.75 | 66.95 ± 8.48 |
| GPT (U/L) | 41.70 ± 19.63 | 25.20 ± 9.81 * |
| AP (U/L) | 24.58 ± 17.86 | 18.77 ± 15.49 |
| Urea (mg/dl) | 143.60 ± 147.93 | 5.97 ± 13.28 * |
| Creatinin (mg/dl) | 0.50 ± 0.17 | 0.41 ± 0.086 |
| Abumin (g/dl) | 2.98 ± 0.11 | 2.90 ± 0.089 |
| Bilirrubin (mg/dl) | 1.28 ± 0.22 | 1.15 ± 0.26 |
| **URINE** | | |
| | ***Coq9*$^{+/+}$** | ***Coq9*$^{+/+}$ + 0.33 % β-RA** |
| | Mean ± SD | |
| Albumin (g/dl) | 0.06 ± 0.026 | 0.03 ± 0.021 |
| Total Protein (g/dl) | 0.80 ± 0.46 | 0.72 ± 0.21 |
| Creatinin (mg/dl) | 31.39 ± 7.71 | 30.40 ± 4.24 |
| Urea (mg/dl) | 78.96 ± 125.65 | 3.72 ± 1.54 |
| Uric Acid (mg/dl) | 1.80 ± 1.70 | 1.72 ± 0.82 |
| Phosphorus (mg/ml) | 66.63 ± 26 | 77.02 ± 18.14 |
| Calcium (mg/dl) | 7.83 ± 4.83 | 6.79 ± 3.07 |
| Magnesium (mg/dl) | 5.47 ± 0.77 | 5.65 ± 1.77 |

#### Figures

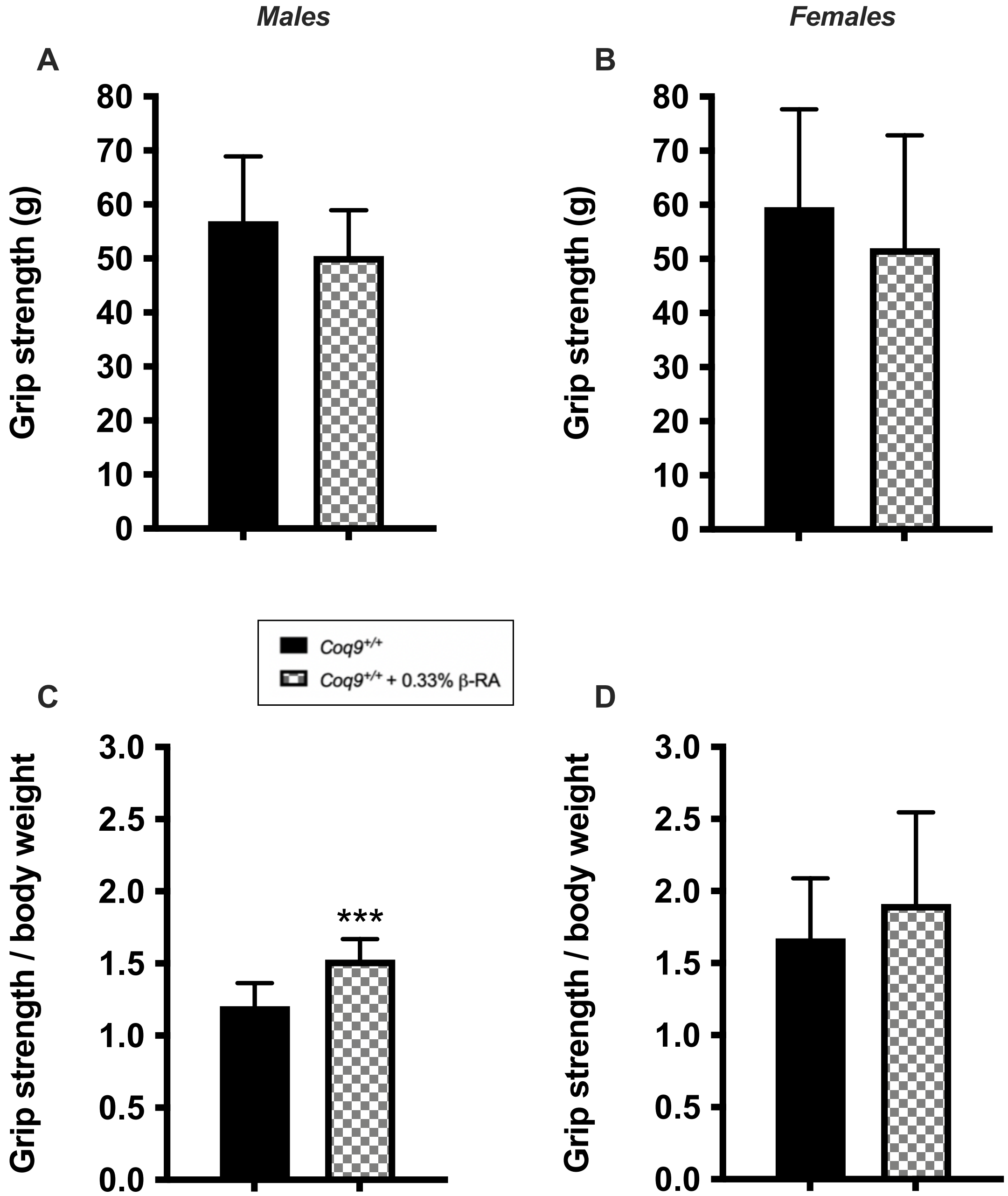


**Figure S1**. **Muscle Strength.**
(**A** to **D)** Grip test in the hind legs of male (A, C) and female (B, D) wild-type mice supplemented with 0.33% β-RA. Data are expressed as grip strength (A, B) and grip strength normalized by the body weight (C, D).
Data are expressed as mean ± SD. ***P < 0.001, differences *versus* *Coq9*$^{+/+}$ (*t*-test; n = 6–9 for each group).

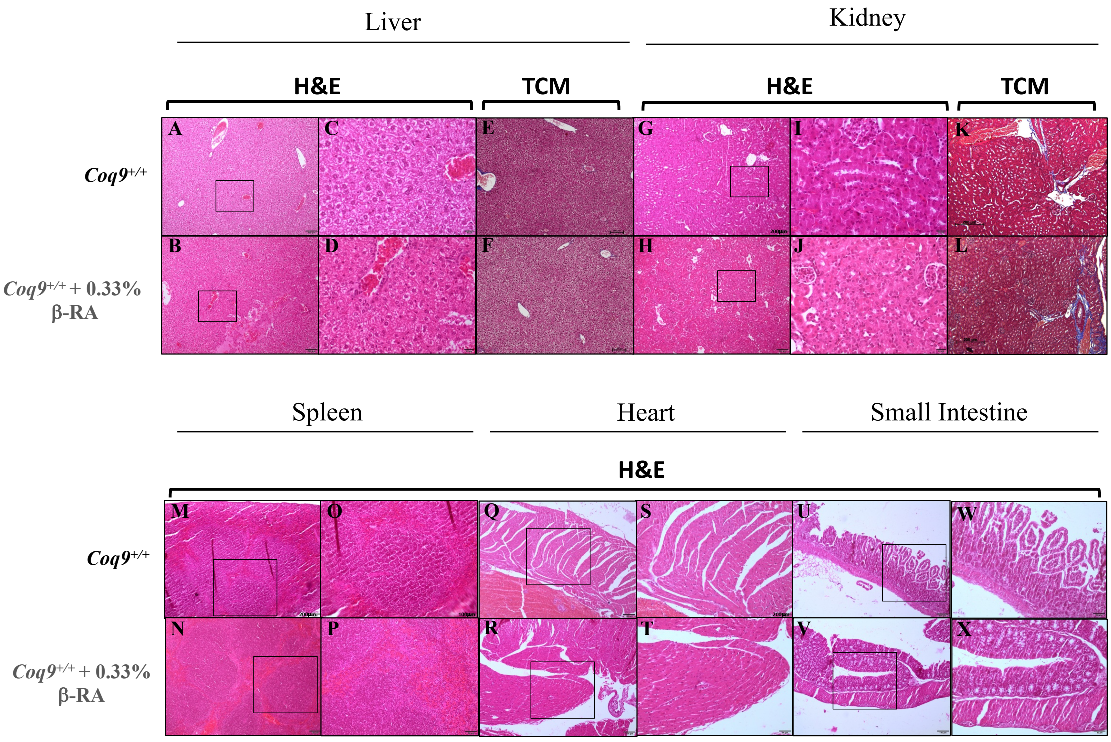


**Figure S2**. **Morphological and histological features of the liver, kidney, spleen, heart and small intestine from *Coq9*$^{+/+}$ *and Coq9*$^{+/+}$ mice under 0.33% of β-RA treatment.**

(**A** to **D)** H&E stain in the liver from *Coq9*$^{+/+}$ mice and *Coq9*$^{+/+}$ mice under 0.33% of β-RA treatment.

(**E** to **F)** Trichrome Gomori (TCM) stain in the liver from *Coq9*$^{+/+}$ mice and *Coq9*$^{+/+}$ mice under 0.33% of β-RA treatment.

**(G** to **J)** H&E stain in the kidney from *Coq9*$^{+/+}$ mice and *Coq9*$^{+/+}$ mice under 0.33% of β-RA treatment.

**(K** to **L)** TCM stain in the kidney from *Coq9*$^{+/+}$ mice and *Coq9*$^{+/+}$ mice under 0.33% of β-RA treatment.

(**M** to **P)** H&E stain in the spleen from *Coq9*$^{+/+}$ mice and *Coq9*$^{+/+}$ mice under 0.33% of β-RA treatment.

(**Q** to **T)** H&E stain in the heart from *Coq9*$^{+/+}$ mice and *Coq9*$^{+/+}$ mice under 0.33% of β-RA treatment.

(**U** to **X)** H&E stain in the gut from *Coq9*$^{+/+}$ mice and *Coq9*$^{+/+}$ mice under 0.33% of β-RA treatment.

Tissues from mice at 3 months of age.

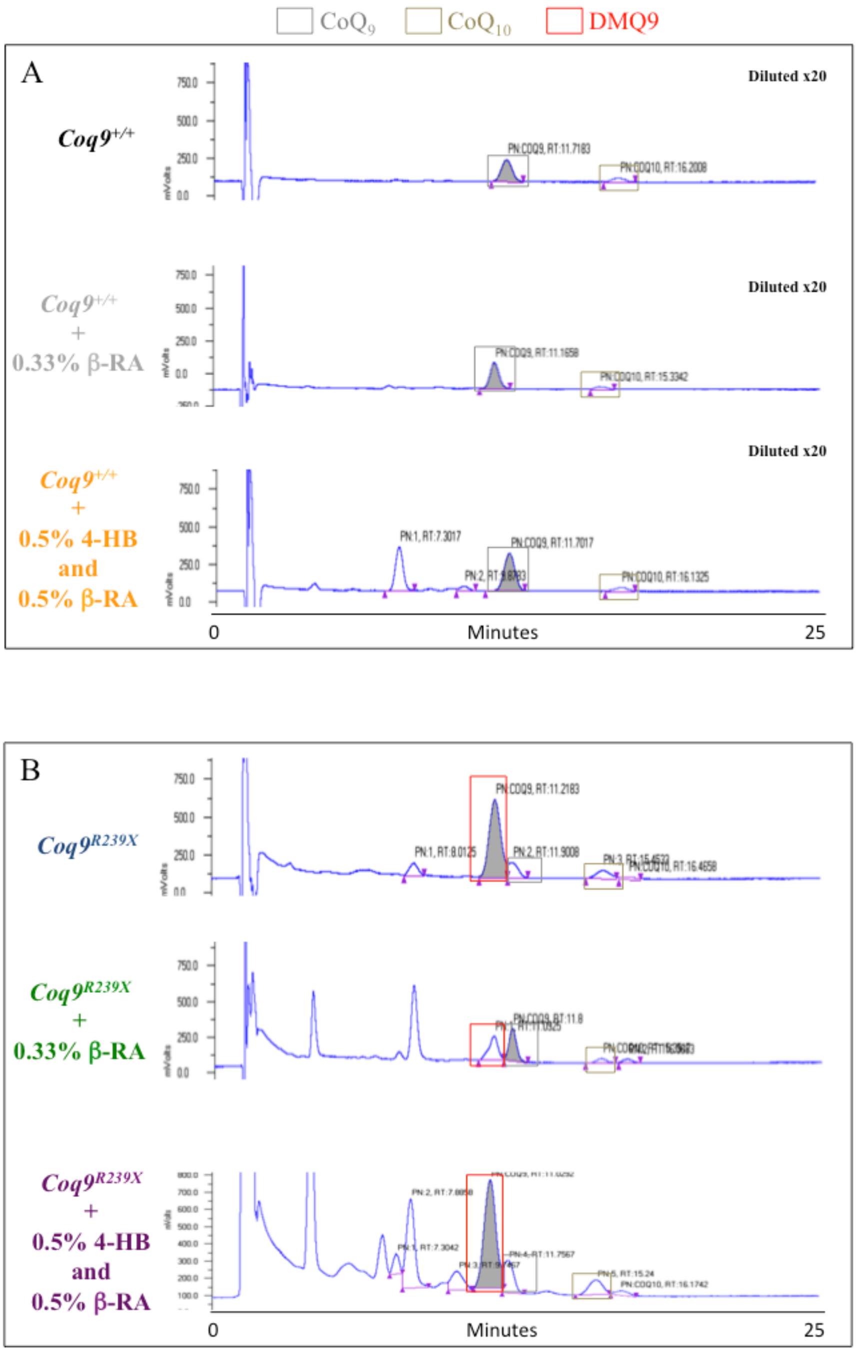


**Figure S3. Representative chromatographs showing the peaks of $CoQ_9$ and $DMQ_9$ in the kidneys.**

(**A**) Chromatographs for $CoQ_9$ in the kidney of a *Coq9*$^{+/+}$ mouse, *Coq9*$^{+/+}$ mouse under 0.33% of β-RA treatment, and *Coq9*$^{+/+}$ mouse under 0.5% of 4-HB + 0.5% of β-RA treatment.

(**B**) Chromatographs for $CoQ_9$ and $DMQ_9$ in the kidney of a *Coq9*$^{R239X}$ mouse, *Coq9*$^{R239X}$ mouse under 0.33% of β-RA treatment, and *Coq9*$^{R239X}$ mouse under 0.5% of 4-HB + 0.5% of β-RA treatment.

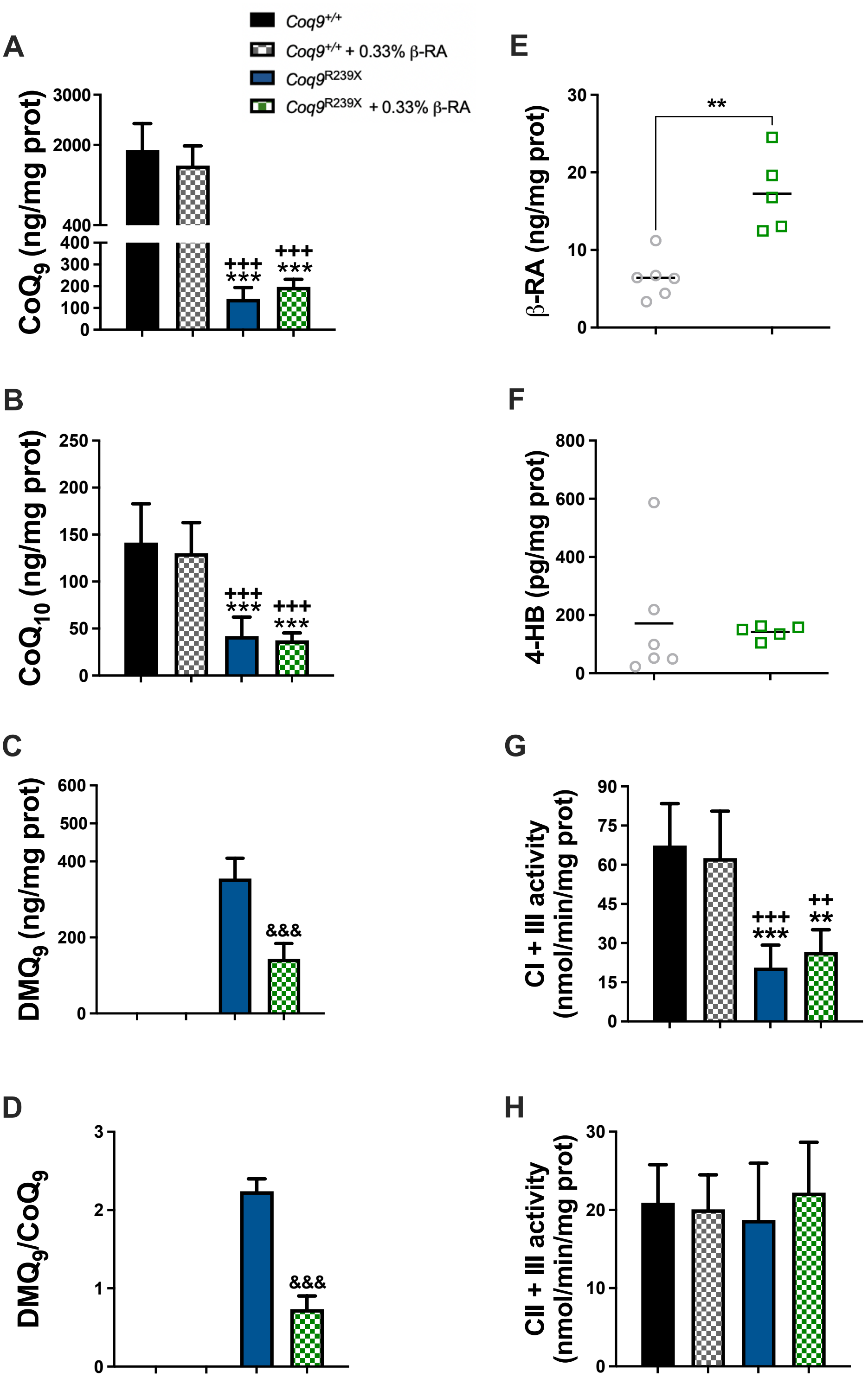


**Figure S4. CoQ metabolism and mitochondrial function in the heart from *Coq9*$^{+/+}$ mice, *Coq9*$^{+/+}$ mice under the supplementation with 0.33% β-RA, *Coq9*$^{R239X}$ mice and *Coq9*$^{R239X}$ mice under the supplementation with 0.33% β-RA.**
(**A**) Levels of $CoQ_9$; (**B**) Levels of $CoQ_{10}$; (**C**) Levels of $DMQ_9$; (**D**) $DMQ_9/CoQ_9$ ratio; (**E**) Levels of β-RA; (**F**) Levels of 4-HB; (**G**) Complex I + III (CI + III) activities; (**H**) Complex II+ III (CII + III) activity.

Tissues from mice at 3 months of age. Data are expressed as mean ± SD. **P < 0.01; ***P < 0.001; differences versus *Coq9*$^{+/+}$. ++P < 0.01; +++P < 0.001, differences *versus* *Coq9*$^{+/+}$ under 0.33% β-RA treatment (one-way ANOVA with a Tukey's post hoc test or t-test; n = 5–8 for each group). Note that $DMQ_9$ is not detected in samples from *Coq9*$^{+/+}$ mice.

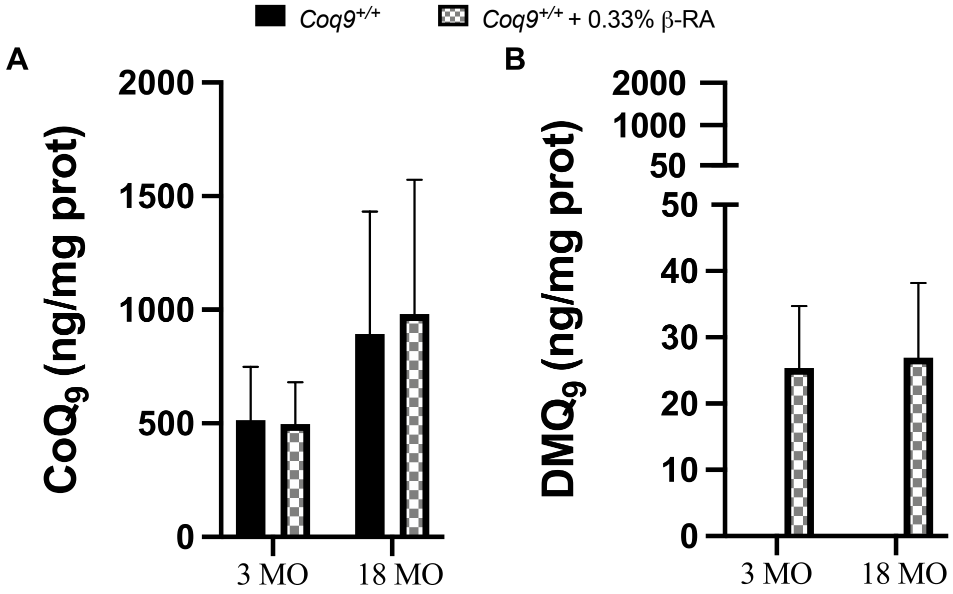


**Figure S5. CoQ levels in WAT from *Coq9*$^{+/+}$ mice *and Coq9*$^{+/+}$ mice under the supplementation with 0.33% β-RA.**
(**A**) Levels of $CoQ_9$; (**B**) Levels of $DMQ_9$.
Tissues from mice at 3 or 18 months of age (MO). Note that $DMQ_9$ is not detected in samples from *Coq9*$^{+/+}$ mice.

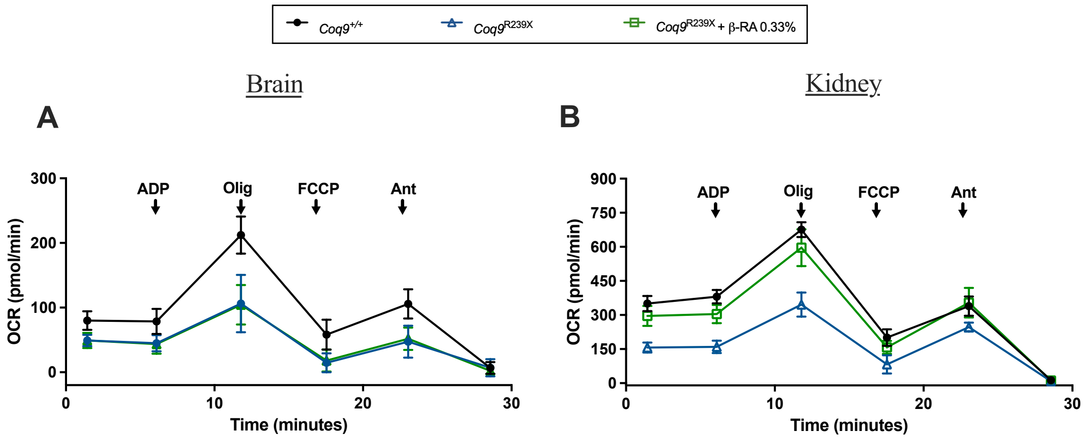


**Figure S6. Mitochondrial oxygen consumption rate (represented as State 3o, in the presence of ADP and substrates) in brain (A) and kidneys (B).** ADP = Adenosine diphosphate; Olig = Oligomycin, a mitochondrial ATP synthase inhibitor; FCCP = Carbonyl cyanide-p-trifluoromethoxyphenylhydrazone, an uncoupler of mitochondrial respiration and ATP synthesis; Ant = Antimycin A, a mitochondrial Complex III inhibitor.

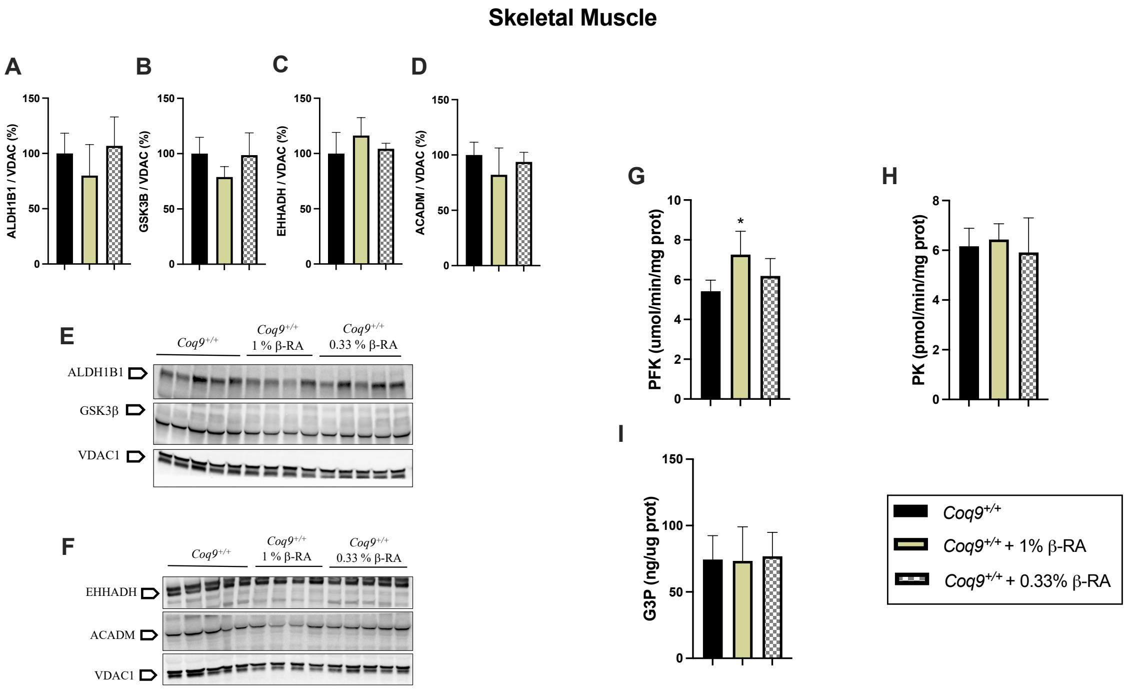


**Figure S7. Metabolic characterization of the skeletal muscle after the treatment with β-RA in *Coq9*$^{+/+}$ mice.**

(**A** to **F**) Levels of the proteins ALDH1B1 (A, E), GSK3β (B, E), EHHADH (C, F) and ACADM (D, F) in the skeletal muscle of *Coq9*$^{+/+}$ mice treated with β-RA at 1% and 0.33%. VDAC1 was used as loading control. The experiments were performed in tissue homogenate.

(**G** to **I**) Activities of the glycolytic enzymes Phosphofructokinase (PFK) (G) and Pyruvate Kinase (PK) (H) in the and skeletal muscle; levels of Glycerol-3-Phosphate (G3P) (I) in the skeletal muscle (S).

Tissues from mice at 3 months of age. Data are expressed as mean ± SD. *P < 0.05; differences versus *Coq9*$^{+/+}$ (one-way ANOVA with a Tukey's post hoc test or t-test; n = 5–7 for each group).

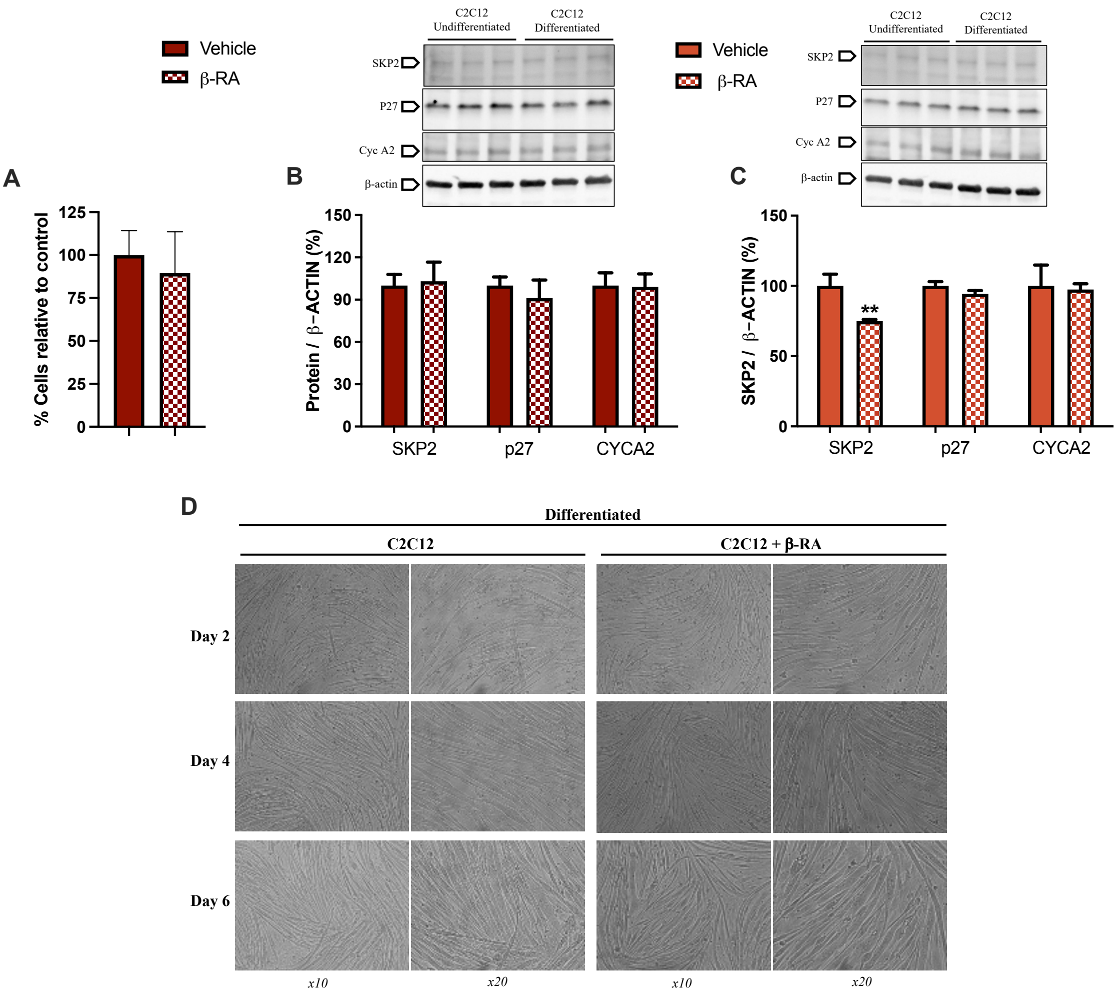


**Figure S8. Effects of β-RA in the proliferation and differentiation of C2C12 myoblasts.**

(**A**) Percentage of C2C12 cells after the treatment with 1mM β-RA, relative to the number of untreated C2C12 cells. Cells cultured in proliferative conditions.

(**B**) Levels of the proteins SKP2, p27 and CYCA2, which are involved in the control of the cell cycle. C2C12 cells were treated for seven days with 1mM β-RA in proliferative conditions.

(**C**) Levels of the proteins SKP2, p27 and CYCA2, which are involved in the control of the cell cycle. C2C12 cells were treated for seven days with 1mM β-RA in differentiative conditions.

(**D**) Images of the C2C12 cells under the microscope in differentiative conditions. C2C12 cells were treated with 1mM β-RA and the images were taken in different days (2, 4 and 6).

Data are expressed as mean ± SD. **P < 0.01, differences *versus* untreated cells (t-test; n = 6 for each group).

**WAT**

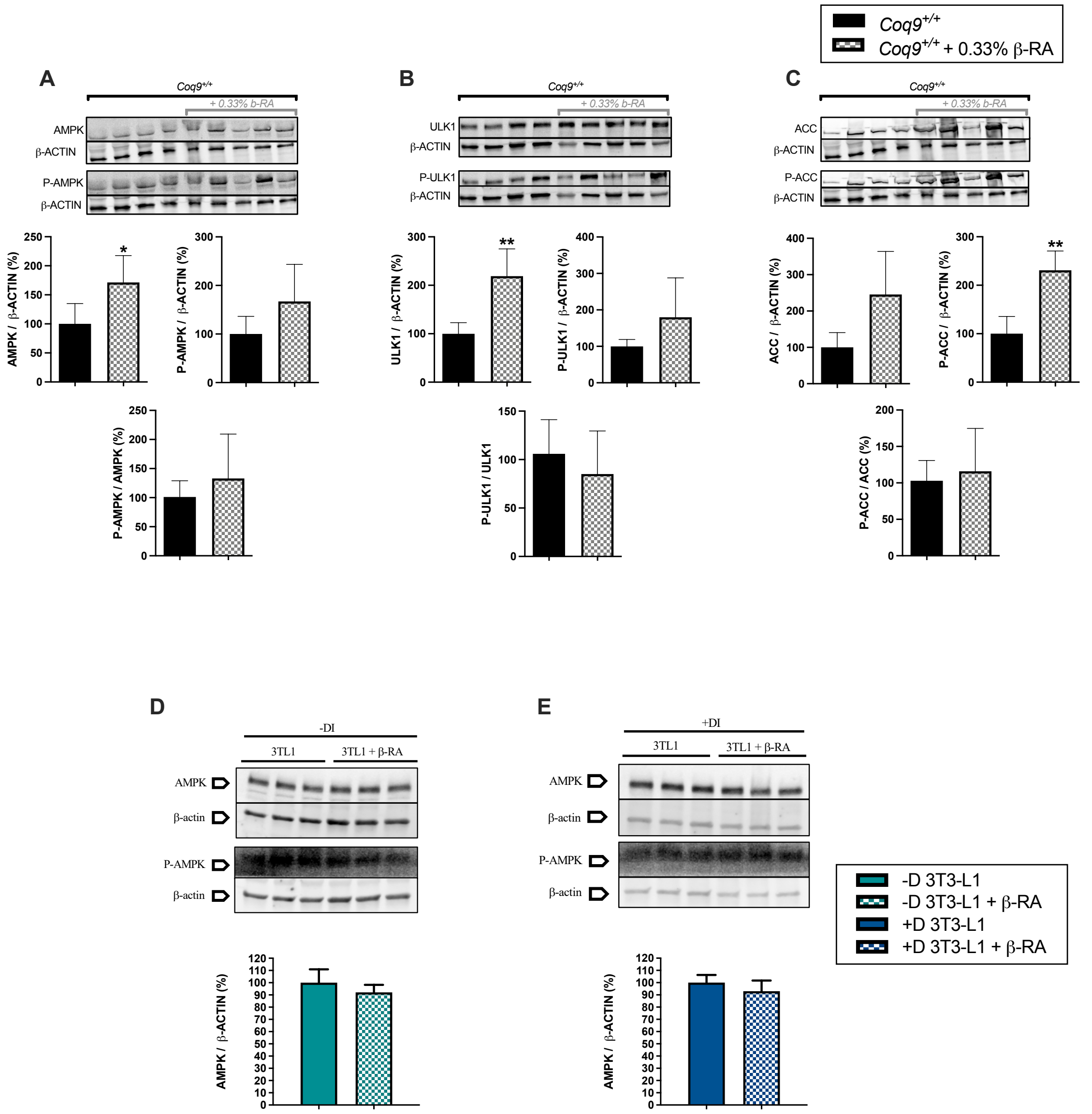


**Figure S9. Analysis of the AMPK pathway in white adipose tissues and 3TL1 cells.**
(**A**) Levels of AMPK and p-AMPK, and p-AMPK/AMPK ratio in epididymal WAT from β-RA-treated and untreated wild-type mice.
(**B**) Levels of ULK1 and p-ULK1, and p-ULK1/ULK1 ratio in epididymal WAT from β-RA-treated and untreated wild-type mice.
(**C**) Levels of ACC and p-ACC, and p-ACC/ACC ratio in epididymal WAT from β-RA-treated and untreated wild-type mice.
(**D**) Levels of AMPK and p-AMPK, and p-AMPK/AMPK ratio in 3TL1 cells treated with 1mM β-RA in proliferative conditions for seven days.
(**E**) Levels of AMPK and p-AMPK, and p-AMPK/AMPK ratio in 3TL1 cells treated with 1mM β-RA in differentiative conditions for seven days.
Tissues from mice at 3 months of age (A to C). Data are expressed as mean ± SD. *P < 0.05; **P < 0.01, differences *versus* *Coq9*$^{+/+}$ mice (t-test; n = 4-6 for each group).

#### Legends to Supplementary Videos

**Movie S1.** Video that shows the difference between a *Coq9*$^{+/+}$ mouse and a *Coq9*$^{R239X}$ mouse under 0.33% β-RA supplementation, both males at 20 months of age. Both animals have a healthy appearance, although the treated *Coq9*$^{R239X}$ mouse is smaller, as previously reported.

**Movie S2.** Video that shows the difference between a *Coq9*$^{R239X}$ mouse and a *Coq9*$^{R239X}$ mouse under 0.33% β-RA treatment, both males at 3 months of age. The untreated *Coq9*$^{R239X}$ mouse has developed a paralysis in the legs, although the treated *Coq9*$^{R239X}$ mouse has a healthy appearance.

**Movie S3.** Video that shows a *Coq9*$^{+/+}$ mouse and a *Coq9*$^{+/+}$ mouse under 0.33% β-RA supplementation, both males at 20 months of age. The appearance of both animals is similar.
